# An oligarchy of NO-producing interneurons controls basal and evoked blood flow in the cortex

**DOI:** 10.1101/555151

**Authors:** Christina T. Echagarruga, Kyle Gheres, Patrick J. Drew

## Abstract

Changes in cortical neural activity are coupled to changes in local arterial diameter and blood flow. However, the neuronal types and the signaling mechanisms that control the basal diameter of cerebral arteries or their evoked dilations are not well understood. Using chronic two-photon microscopy, electrophysiology, chemogenetics, and pharmacology in awake, head-fixed mice, we dissected the cellular mechanisms controlling the basal diameter and evoked dilation in cortical arteries. We found that modulation of overall neural activity up or down caused corresponding increases or decreases in basal arterial diameter. Surprisingly, modulation of pyramidal neuron activity had minimal effects on basal or evoked arterial dilation. Instead, the neurally-mediated component of arterial dilation was largely regulated through nitric oxide released by neuronal nitric oxide synthase (nNOS)-expressing neurons, whose activity was not reflected in electrophysiological measures of population activity. Our results show that cortical hemodynamic signals are not controlled by the average activity of the neural population, but rather the activity of a small ‘oligarchy’ of neurons.

## Introduction

The brain is an energetically demanding organ, and understanding the mechanisms by which neural activity regulates local blood flow is a fundamental issue in neuroscience. Changes in neural activity are coupled to blood flow via neurovascular coupling, where neural activity drives the dilation of arterioles and other vessels^1–4^. Typically, increases in the gamma-band power of the LFP (local field potential), a reliable indicator of overall neural activity, are correlated with vasodilation in the awake animal^5–8^. However, under some conditions, neural activity (as measured with electrophysiological recordings) and hemodynamic signals are decoupled^7, 9–11^. There are multiple signaling pathways implicated in linking neural activity, both directly and indirectly, to arteriole dilations^12, 13^. Whether these dilations are controlled by the activity of excitatory pyramidal neurons^14–16^ or interneurons^17–19^ remains unresolved. Elucidating which neuronal types control arterial dilation^20^ is important for interpreting and decoding neural activity^21, 22^ from hemodynamic imaging signals such as those used by fMRI^5, 23^. Moreover, the regulation of *basal* arterial diameter, important in controlling baseline blood flow, remains poorly understood. Changes in baseline flow can radically alter the sign and magnitude of evoked BOLD fMRI signals^24^, which depend on the ratio of the change in blood flow to the baseline. Additionally, decreases in basal cerebral blood flow occur with aging, and the amplitude of these decreases predicts the onset of neurodegeneration^25^, which implies adequate basal blood flow is important for a healthy brain. Thus, understanding the neural control of both basal and evoked changes in arterial diameter is essential for interpreting functional imaging and understanding the pathophysiology of disease. Here, we show that the activity of a small group of neurons that express nNOS and the substance P receptor (NK1R) are the predominant regulators of cortical hemodynamic signals, and whose activity is invisible to traditional extracellular recordings of neural activity.

### Basal arterial diameter and evoked dilation are controlled by local neural activity

We used two-photon microscopy^26^ to chronically image pial and penetrating arterioles in the somatosensory cortex of awake, head-fixed mice (Fig. 1A and B)^8, 27^ through polished and reinforced thinned-skull (PoRTs) windows^28^. Because we were able to repeatedly image from the same vessels (Fig. 1C), we were able to make quantitative comparisons of the responses of individual vessel to experimental modulations in neural activity. As we measured multiple vessels from each mouse, we used linear mixed-effect models (see Methods)^29^ to account for within-animal correlations^7, 30^ that make methods such as ANOVA inappropriate^29^. Using cytochrome oxidase staining, we reconstructed the locations of individual arterioles relative to the forelimb representation in somatosensory cortex (purple and green shaded regions Fig. 1B, top) that is strongly activated by locomotion. We measured arteriole diameter dynamics driven by locomotion, as behavior is the primary driver of hemodynamic signals in the awake brain (Fig. 1D)^7, 27^, and because sensory stimulation in awake animals invariably elicits movement^31, 32^. During bouts of voluntary locomotion, neural activity increased substantially in the forelimb/hindlimb (FL/HL) representation in somatosensory cortex, as measured by changes in firing rate or power in the gamma-band of the LFP (Fig. S1)^33^. In response to this increase in neural activity, both pial and penetrating arterioles in the FL/HL representation dilated^27, 34^ (Fig. 1D and Fig. S2A and Supplementary movies 1 & 2) with short latency (onset time: 0.59±0.31 seconds in the FL/HL region).

**Figure 1.**
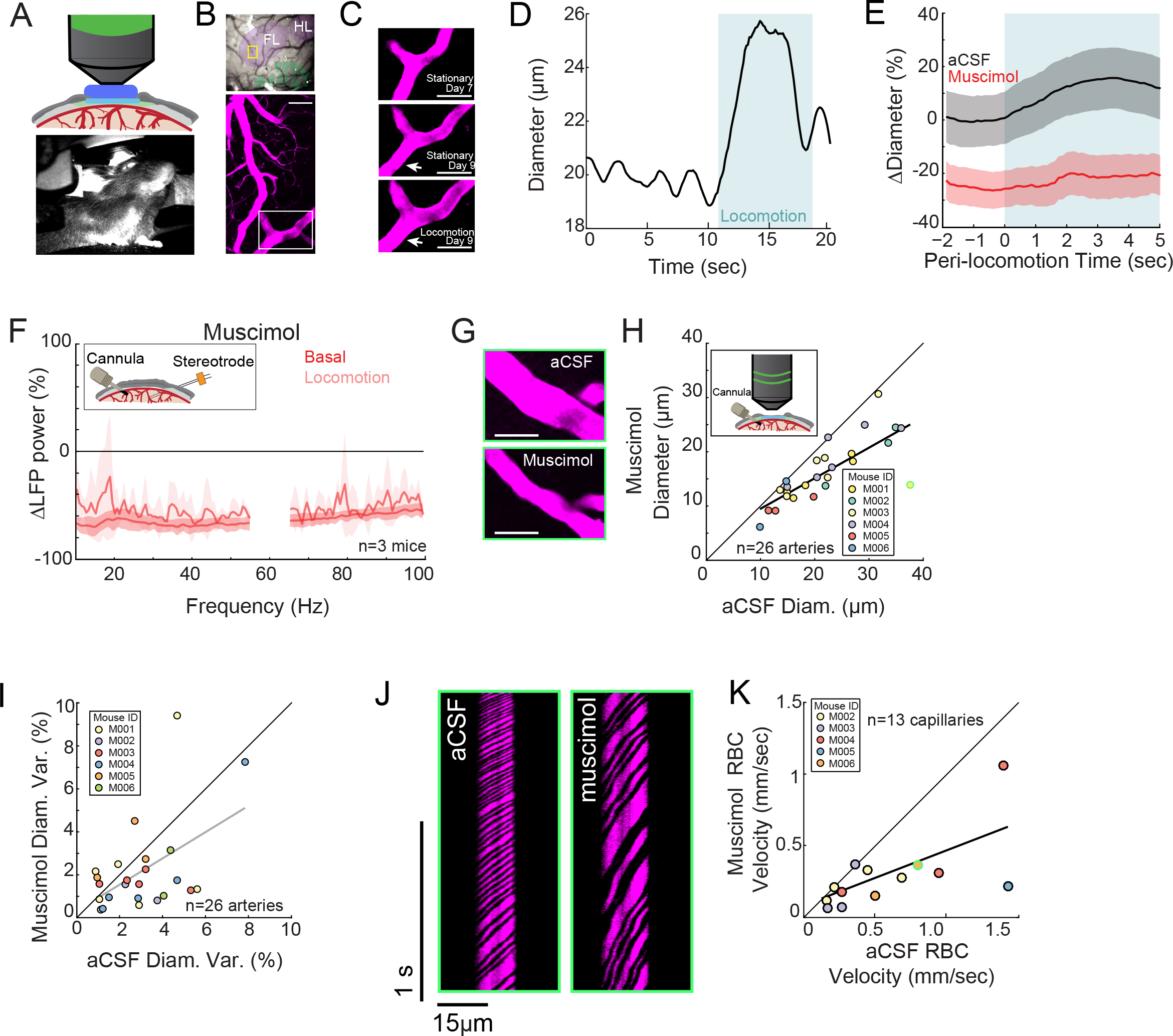
Local neural activity controls basal and evoked arteriole diameter. **A.** Top, schematic of imaging window. Bottom, photo of mouse on spherical treadmill. **B.** Top, photo of the pial vasculature of the somatosensory cortex through the PoRTs window. Cytochrome oxidase processing was used to localize the forelimb/hindlimb (FL/HL, purple) and vibrissae cortex (green). Bottom, a maximum projection of two-photon images of vasculature (scale bar 50μm) within the yellow box in the top image. **C.** Top and middle, two-photon images (from white box in B, bottom) of the same arteriole 7 and 9 days after window implantation (scale bars 50μm), both taken when mouse was stationary. The bottom image shows an example image 9 days after implantation during locomotion (scale bar 50μm) **D.** Locomotion induces rapid dilation in pial arterioles. Diameter (region marked by white arrow in **C**) was plotted versus time. Blue shading denotes period of locomotion. **E.** Population locomotion-triggered averages following aCSF (black) and muscimol (red) infusions (n=6 mice, 12 arterioles in FL/HL representation). For both cases, the individual diameters are normalized by the average basal diameter of the vessel after vehicle infusion. Note the rapid rise to peak in the control, and the lack of dilation in muscimol-infused mice, showing the locomotion-triggered response is under local neural control. Shading represents mean ± standard deviation. **F.** LFP power spectra during stationary periods (basal) and locomotion after muscimol infusion, normalized to vehicle infusion in the same animal. Muscimol caused a substantial decrease in gamma band power (basal, −60.9±13.3%, paired t-test p=2.8×10^−3^; locomotion, −52.7±5.3%, paired t-test p=0.01, n=3 mice). Inset, schematic of electrode-cannula preparation **G.** Representative images of a pial arteriole during times of no locomotion after vehicle infusion (top) and after muscimol infusion (bottom). Arteriole diameter decreased after muscimol infusion (scale bar 50μm) **H.** Basal arteriole diameter following vehicle infusion plotted versus basal diameter after muscimol infusion. Each point is a single vessel, and the mouse identity is represented by the color. The black line shows the linear regression of aCSF vs. muscimol diameter, and muscimol caused a significant decrease in diameter (−25.4±2.3%, LME p=3.3×10^−4^, n=6 mice, 26 vessels). Insets show schematic of PoRTs window-cannula preparation and mouse ID color code. The point outlined in green is representative image from G. **I.** Scatter plot of basal arteriole diameter variance during stationary periods following vehicle infusion (x-axis) versus muscimol infusion (y-axis). There was no significant change in arterial diameter (−14.2±67%, LME p=0.06, n=6 mice, 26 vessels). The variance measurements are from the same vessels measured in H. **J.** Representative space-time image of linescans of the same capillary after aCSF or muscimol infusion. **K.** Basal red blood cell (RBC) velocity plotted after vehicle infusion (x-axis) vs after muscimol infusion (y-axis). Muscimol caused a significant decrease in the RBC velocity (−44±27%, LME p=0.03, n=5 mice, 13 capillaries). The point outlined in green is the same vessel that the linescans in J are taken from.

We first determined the role of overall local neural activity in maintaining baseline arterial diameter and for mediating hemodynamic changes during locomotion. We suppressed the activity of all neurons via intracranial infusions of muscimol (a GABAA receptor agonist) (Fig. 1F), and compared the changes in neural activity and arterial diameter dynamics to artificial cerebrospinal fluid (aCSF) infusions in the same mouse. Infusions of muscimol substantially decrease neural activity ~1.5mm from the cannula^7^, and so will suppress activity in all cortical layers, unlike topical administration of drugs which only affect the supragranular layers^35^. Cannula implantations do not alter hemodynamic responses^7^. We found that after the infusion of muscimol, there was a 60.9±1.0% decrease in baseline gamma-band power (Fig. 1F, line, paired t-test p=2.8×10^−3^) and 52.7±5.3% decrease in the gamma-band power during locomotion relative to aCSF infusions (Fig. 1F, paired t-test p=0.01). We found that muscimol infusions led to a substantial decrease in the basal arterial diameter (−25.4±2.3%, Figure 1G-H, LME p=3.3×10^−4^). We refer to the diameter of the vessel during periods lacking locomotion (see Methods) as the “basal” diameter. Note that the locomotion-triggered averages calculated using the subset of vessels histologically identified to be in the FL/HL representation, while the changes in basal diameters were calculated using all vessels imaged in the somatosensory cortex. The effects of muscimol infusions were not due to direct actions of the cerebral vasculature, which lack GABAA receptors36,37, but rather by the removal of tonic vasodilatory signals released by neurons. The locomotion-evoked response was nearly completely blocked by local silencing of neurons, showing that these arterial dilations were not due to non-specific cardiovascular responses^38^ or modulatory input from other brain regions directly onto the vessels^39^, but rather controlled by local activity. Similar changes were observed in penetrating arterioles (Fig. S2B and C), and mice continued to spontaneously locomote after infusions of muscimol and other manipulations of neural activity (Fig. S3). Suppressing neural activity with muscimol did not abolish spontaneous fluctuations in arterial diameter (Fig. 1I), consistent with previous reports of autonomous oscillations in arterioles^7^. To determine if this decrease in arterial basal diameter drove a decrease in blood flow, we then measured red blood cell (RBC) velocity in the capillaries^40^ (Fig. 1J and 1K). Capillary RBC velocity was decreased in muscimol-infused mice relative to aCSF infusions (−44±27%, n=13 capillaries, LME p=0.03, Fig 1K). We observed no significant change in the diameter of the capillaries following the infusion of muscimol (0.12±1.1μm, LME p=0.66, Fig. S4B). Taken together, these results show that baseline levels of neural activity have a tonic vasodilatory effect on cerebral arteries, and decreasing neural activity causes a corresponding decrease in arterial diameter and blood flow.

### Bi-directional chemogenetic manipulation of local neural activity controls basal arterial diameter

We then asked if we could bi-directionally manipulate arterial diameters using chemogenetic techniques to drive changes in overall neural activity. We expressed either hM4D-G(i) or hM3D-G(q)-coupled DREADDs^41^ pan-neuronally under a synapsin promoter in C57-BL6J mice. We targeted the deeper cortical layers, as there is evidence that hemodynamic responses initiate there^3^ before the dilation propagates to the pial arteries. We imaged arterioles in the DREADD-expressing region of cortex (identified by mCherry expression seen in vivo, Fig. 2A, and histologically, Fig. 2B). Injections of CNO in mice with pan-neuronal expression of hM4D-G(i) DREADDs resulted in a decrease in gamma-band power relative to vehicle injections (CNO gamma-band power −8.7±3.9% relative to vehicle, paired t-test p= 0.18 for basal periods, Fig. 2D). Consistent with the results of pharmacological infusions of muscimol, the decreases in overall neural activity drove a decrease in basal arterial diameter (−7.1±7.0%, Figure 2E, LME p=0.02) and in the locomotion-evoked response (Fig. 2F, Fig. S5A and S5E). In contrast, mice expressing hM3D-G(q) DREADDs, which increases excitability pan-neuronally, showed increases in gamma-band power (+8.46±7.6%, paired t-test p= 2.7×10^−3^ for basal periods, Fig. 2G) with CNO injection relative to vehicle. The basal arterial diameter was also increased when the excitability of all neurons was significantly increased (+18.8±5.6%, LME p=0.03, Fig. 2H). The increase in basal arteriole diameter by hM3D-G(q) DREADD activation was large enough to almost completely occlude the locomotion-evoked response (Fig. 2I). Control mice infected with an AAV expressing only a fluorescent reporter protein did not show significant changes in neural activity or basal arterial diameter after the injection of CNO (Fig. S6B-D), showing CNO has minimal off-target effects in our system. These results demonstrate that neurons, or a subset of neurons, tonically release vasodilator(s), and that the manipulation of this population of neurons up or down increases or decreases the basal arteriole diameter correspondingly.

**Figure 2.**
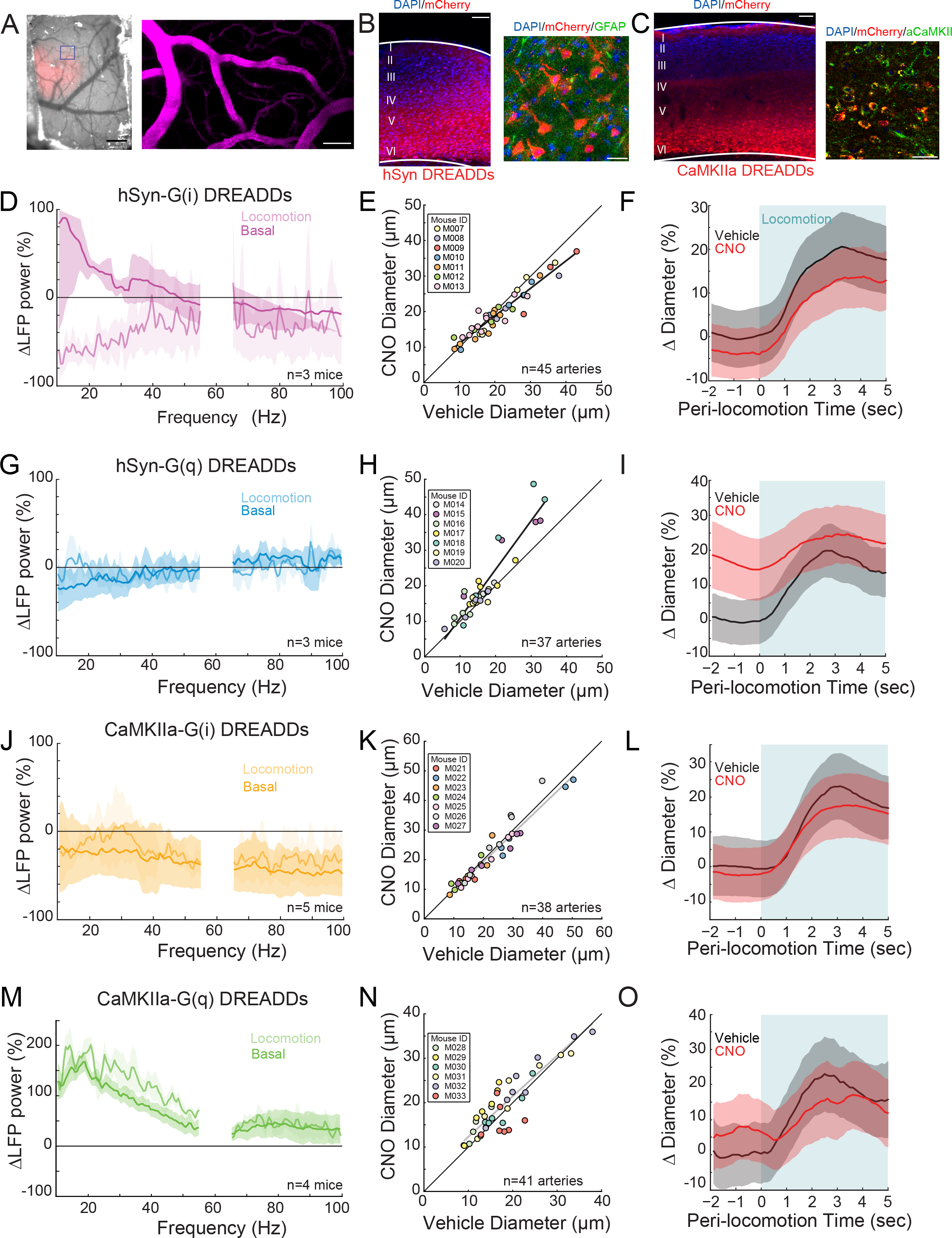
Activity, but not of pyramidal neurons, bidirectionally controls basal arteriole diameter. **A.** Left, image through the polished and reinforced thinned-skull window showing AAV expression (red) in the somatosensory cortex (scale bar 1 mm). Right, two-photon image of vasculature (scale bar 50μm) taken from the purple box on left. **B.** Representative image of AAV-hSYN-HA-hM3D(Gq)-mCherry infected cortex. Left image is wide field image of hSyn-mCherry DREADDs virus (red) in cortex with DAPI (blue) staining (scale bar 100μm). Right is magnified image of hSyn-mCherry DREADDs virus (red) in cortex with DAPI (blue) and GFAP (green) staining (scale bar 30μm), showing the virus was not expressed in astrocytes. Tissues sectioned to a 90μm thickness **C.** Representative image of AAV-CaMKIIa-hM3D(Gq)-mCherry infected cortex, where DREADDs are expressed under a CaMKIIa promoter. Left image is wide field image of CaMKIIa-mCherry DREADDs virus (red) in cortex with DAPI (blue) staining (scale bar 100μm). Right is a magnified image of CaMKIIa-mCherry DREADDs virus (red) in cortex with DAPI (blue) and αCaMKII (green) staining (scale bar 30μm) showing expression in excitatory neurons. Data in **D-F** are from mice injected with AAV-hSYN-HA-hM4D(Gi)-mCherry, data in **G-I** is from mice injected with AAV-hSYN-HA-hM3D(Gq)-mCherry. **D.** LFP power spectra during stationary periods (basal) and locomotion after CNO injection in hSyn G(i) DREADDs mice, normalized to vehicle injection in the same animal. Pan-neuronal expression of G(i) DREADDs reduces neural activity in the gamma band (basal, −8.7±3.9%, paired t-test p= 0.18; locomotion, −29.1±37 %, paired t-test p=0.38, n=3). Shading denotes mean ± standard deviation. **E.** Plot of basal arteriole diameter after vehicle injection (x-axis) versus CNO injection (y-axis). CNO caused a significant decrease in arterial diameter (−7.1±7.0%, LME p=0.02 Bonferroni corrected, n=7 mice, 45 vessels). **F.** Population locomotion-triggered averages after vehicle (black) and CNO (red) injections (n=7 mice, 33 arterioles in FL/HL representation). For both cases, the diameters were normalized by the average basal diameter of the vessel after vehicle injection. **G.** LFP power spectra during stationary periods (basal) and locomotion after CNO injection in hSyn G(q) DREADDs mice, normalized to vehicle injection of the same mouse. Pan-neuronal expression of G(q) DREADDs increases neural activity in the gamma band (basal, 8.46±7.6%, paired t-test p= 2.7×10^−3^; locomotion, +0.2±10.3 %, paired t-test p=0.21, n= 3). **H.** Plot of basal arteriole diameter after vehicle injection (x-axis) versus CNO injection (y-axis). CNO caused a significant increase in arterial diameter (+18.8±5.6%, LME p=0.03 Bonferroni corrected, n= 7 mice, 37 vessels). **I.** Population locomotion-triggered averages in response to locomotion after vehicle (black) and CNO (red) injections (n= 7 mice, 25 arterioles in FL/HL representation). For both cases, the diameters were normalized by the average basal diameter of the vessel after vehicle injection. Increasing neural activity caused a basal arterial dilation that nearly completely occluded the locomotion-triggered response. Data in **J-L** are from mice injected with AAV-CaMKIIa-hM4D(Gi)-mCherry, data in **M-O** is from mice injected with AAV-CaMKIIa-hM4D(Gq)-mCherry (expression in pyramidal neurons) **J.** LFP power spectra during stationary periods (basal) and locomotion after CNO injection in CaMKIIa G(i) DREADDs mice, normalized to vehicle injection of the same mouse. Pyramidal neuron expression of G(i) DREADDs reduces neural activity in the gamma band (basal, −40.26±3.3%, paired t-test p=0.05; locomotion, −30.9±18.1 %, paired t-test p=0.019, n=5). **K.** Plot of basal arteriole diameter after vehicle injection (x-axis) versus CNO injection (y-axis). There was no significant change in the diameter (−2.2±5.7 %, LME p=1 Bonferroni corrected, n=7 mice, 38 arterioles). **L.** Population locomotion-triggered averages after vehicle (black) and CNO (red) injections (n=7 mice, 33 arterioles in FL/HL representation). For both cases, the diameters were normalized by the average basal diameter of the vessel after vehicle injection. **M.** LFP power spectra during stationary (basal) and locomotion periods after CNO injection in CaMKIIa G(q) DREADDs mice, normalized to vehicle injection of the same mouse. Pyramidal neuron expression of G(q) DREADDs increased neural activity in the gamma band (basal, +41.23±18.7%, paired t-test p=0.03; locomotion, +54.9±53.3 %, paired t-test p=4.4×10^−3^, n=4). **N.** Plot of basal arteriole diameter after vehicle injection (x-axis) versus CNO injection (y-axis). There was no significant change in diameter (+7.2±8.8%, LME p=0.66 Bonferroni corrected, n=6 mice, 41 vessels). **O.** Population locomotion-triggered averages after vehicle (black) and CNO (red) injections (n=6 mice, 33 arterioles in FL/HL representation). For both cases, the diameters were normalized by the average basal diameter of the vessel after vehicle injection.

### Pyramidal neural activity drives large population neural activity changes without corresponding changes in arterial diameter

The importance of local neural activity on the regulation of the vasculature raises the question of which sub-population(s) of neurons are involved in the signaling. We sought to manipulate the activity of pyramidal neurons by injecting AAVs encoding hM3D(q) and hM4D(i) DREADDs under a CaMKIIa promoter, restricting the expression of the DREADDs to pyramidal cells^42^. CaMKIIa-DREADDs labeled cells colocalized with CaMKIIa expression, and are primarily localized to layer 5 (Fig. 2C). Previous work has shown that optogenetic stimulation of layer 5 pyramidal neurons drives increases in neural activity in all layers, similar to what is seen with sensory stimulation^16, 43^. The dilatory signal in the deeper cortical layers is electrically conducted through the vasculature by gap junctions to the pial vessels^3^. Expressing DREADDs in excitatory neurons produced large changes in overall neural activity as measured with extracellular electrophysiology. Activating hM4D(i) DREADDs expressed in excitatory neurons with CNO caused large decreases in basal gamma-band power (CNO gamma-band power −40.26±3.3% relative to vehicle, Fig. 2J, paired t-test p=0.05). When hM3D(q) DREADD receptors in excitatory neurons were activated, a corresponding increase in basal gamma-band power was observed (CNO gamma-band power increased by 41.23±18.7% relative to vehicle controls, Fig. 2M, paired t-test p=0.03). Despite the large changes in neural activity, we did not see significant changes in the basal arterial diameters with manipulation of pyramidal neuron activity (−2.2±5.7%, LME p=1 for inhibition and +7.2±8.8%, LME p=0.66 for excitation) (Fig. 2K and Fig. 2N, respectively). Note that decreases in basal neural activity generated by activation G(i)-coupled DREADDs in pyramidal neurons were nearly as large as those elicited by muscimol (Fig. S7A), and the increases in basal neural activity generated by activation G(q)-coupled DREADDs in pyramidal neurons were as large as those seen during locomotion (Fig. S7B), yet in neither of these cases did we observe arterial diameter change of similar magnitude.

Previous reports have suggested that the activation of inwardly-rectifying potassium channels by increases in extracellular K^+^ arising from neural activity could mediate vasodilation^44^. To test the role of Kir2.1 channels, we performed experiments where we infused either BaCl2 or ML-133, blockers of Kir2.1 channels. Blocking Kir2 channels with BaCl2 increased basal arterial diameter (+12.49.1±17.2% LME p=6.9×10^−3^, Fig. S8C), but also increased neural activity and induced epileptic-like activity (Fig. S8A), consistent with previous reports of BaCl2 increasing neural excitability45 associated with the blockade of neural K^+^ channels. We obtained similar results with the Kir blocker ML-133^46^, which drove elevations of basal neural activity and basal arterial diameter (+11.8±17.0% LME p=1.3×10^−3^, Fig. S8E and F) which occluded the sensory-evoked response (Fig. S8G). Given the large effects of Kir-channel blockers on neural activity (see also^45^), it is hard to untangle the role of K^+^ sensing on the endothelial cells or smooth muscle directly from any effects caused by increases in overall neural activity.

These results show that arteriole diameter can be completely uncoupled with the standard measures of cortical neuronal activity, which largely reflects activity of excitatory neurons. It also suggests that the activity of the neurons that control local arterial diameter receive minimal input from local pyramidal neurons.

### NO released by nNOS expressing neurons controls arterial diameter independent of overall neural activity

Previous work has implicated neuronally-generated nitric oxide (NO) as a messenger coupling neural activity to arterial vasodilation^4^, though it has also been proposed that NO only modulates the functional hyperemic response^47^. To test the role of neuronal nitric oxide synthase (nNOS) expressing neurons, we expressed hM3D(q) and hM4D(i) DREADDs in nNOS-expressing neurons using flexed AAVs and cre-nNOS mice (B6.129-*Nos1*^*tm1(cre)Mgmj*^/J, Jackson Laboratory #017526). The reporter protein was primarily expressed by a few inhibitory neurons in the deep, infragranular layers (Fig. 3M). Neither suppressing (CNO gamma-band power, −0.80±2.1% relative to vehicle, paired t-test p=0.98, Fig. 3A) nor increasing (CNO gamma-band power −20.17±3.2% relative to vehicle, paired t-test p=0.057, Fig. 3D) the excitability of nNOS-expressing neurons resulted in a significant change in gamma-band power. However, while decreasing the excitability of nNOS neurons did not affect overall neural activity, there was a substantial decrease in basal arteriole diameter (−10.3±6.4% LME p=1.2×10^−4^ Fig. 3B) and a large blunting of the locomotion-induced dilation (Fig. 3C). Elevating the excitability of nNOS expressing neurons did not significantly affect basal or evoked arterial diameter (+5.8±6.6%, LME p=0.66, Fig. 3E). The decreases in evoked dilation were not just due to a shift in the baseline (Fig. S5A and I), showing that nNOS neurons play a role in evoked dilations in addition to basal diameter. The activity of nNOS-expressing neurons impacts both basal and evoked dilations, with minimal changes in overall neural activity.

**Figure 3.**
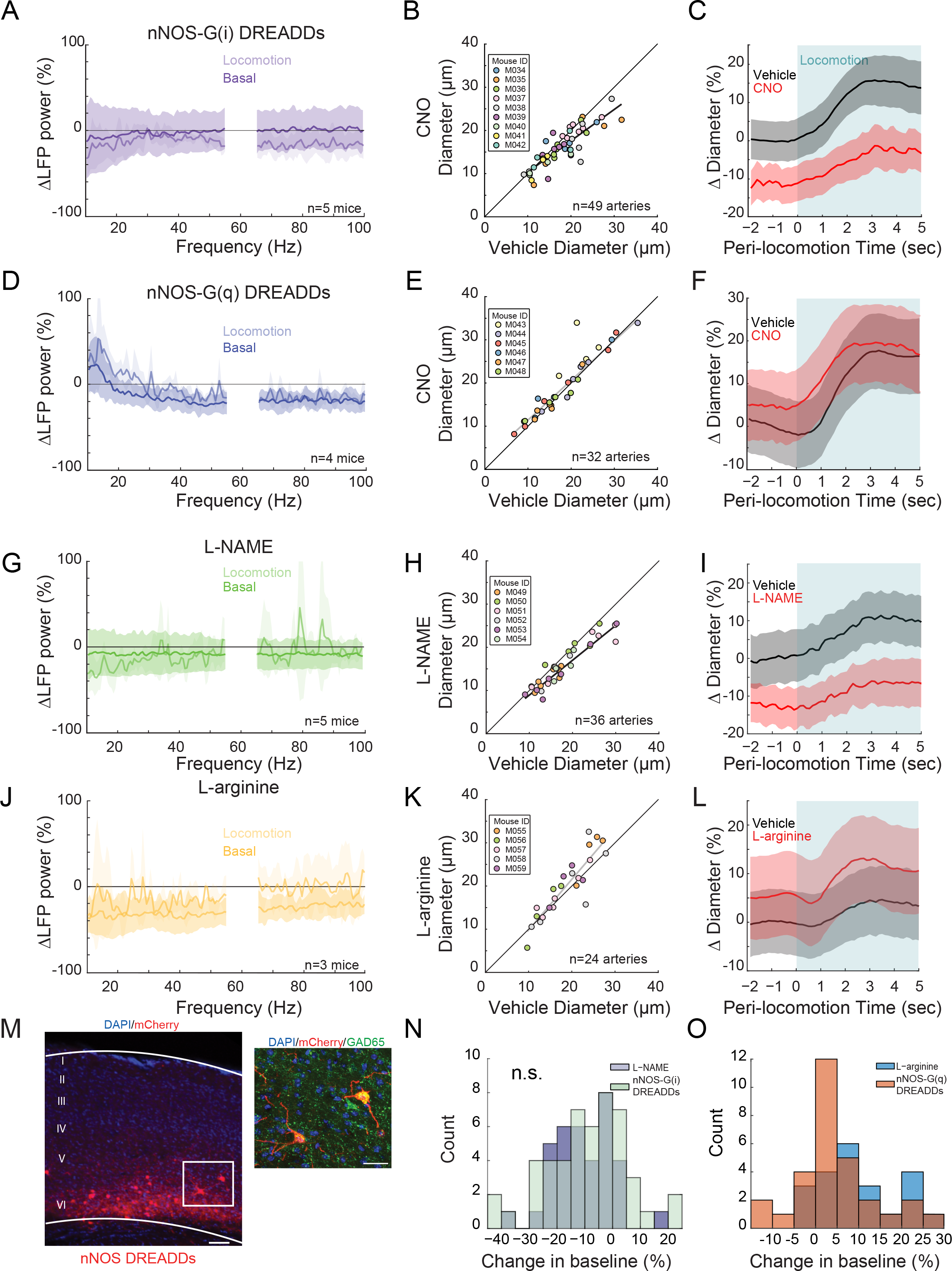
NO released by nNOS expressing neurons controls arteriole diameter independent of overall neural activity. Data in **A-C** are from nNOS-cre mice injected with AAV-hSyn-DIO-hM4D(Gi)-mCherry, data in **D-F** is from nNOS-cre injected with AAV-hSyn-DIO-hM4D(Gq)-mCherry. **A.** LFP power spectra during stationary periods (basal) and locomotion after CNO injection in nNOS G(i) DREADDs mice, normalized to vehicle injection of the same mouse. Expression of G(i) DREADDs in nNOS-positive cells did not change neural activity in the gamma band (basal, −0.80±2.1%, paired t-test p=0.98; locomotion, 13.8±8.2 %, paired t-test p=0.047, n=5). Shading denotes mean ± standard deviation. **B.** Plot of basal arteriole diameter after vehicle injection (x-axis) versus CNO injection (y-axis). CNO caused a significant decrease in the arterial diameter (−10.3±6.4%, LME p=1.3×10^−4^ Bonferroni corrected, n=9 mice, 49 vessels). **C.** Population locomotion-triggered averages after vehicle (black) and CNO (red) injections (n=6 mice, 22 arterioles in FL/HL representation). For both cases, the diameters were normalized by the average basal diameter of the vessel after vehicle injection. Decreasing nNOS+ neural activity reduced the locomotion-triggered dilation. **D.** LFP power spectra during stationary periods (basal) and locomotion after CNO injection in nNOS G(q) DREADDs mice, normalized to vehicle injection of the same mouse. Expression of G(q) DREADDs in nNOS-positive cells reduced neural activity in the gamma band (basal, −20.17±3.2%, paired t-test p=0.057; locomotion, −14.9±8.9 %, paired t-test p= 5.2 x10^−3^, n= 4). **E.** Plot of basal arteriole diameter after vehicle injection (x-axis) versus CNO injection (y-axis). CNO did not cause a significant change in vessel diameters (+5.8±6.6 %, LME p=0.66 Bonferroni corrected, n=6 mice, 32 vessels). **F.** Population locomotion-triggered averages after vehicle (black) and CNO (red) injections (n=6 mice, 18 arterioles in FL/HL representation). For both cases, the diameters were normalized by the average basal diameter of the vessel after vehicle injection. Data in **G-I** are from mice after L-NAME infusions, data in **J-L** is from mice L-arginine infusions. **G.** LFP power spectra during stationary (basal) and locomotion periods after L-NAME infusion, normalized to vehicle infusion in the same mouse. L-NAME has a small, non-significant change neural activity in the gamma band (basal, 8.24±2.8%, paired t-test p=0.39; locomotion, −4.3±10.7%, paired t-test p=0.67, n=5). **H.** Plot of basal arteriole diameter after vehicle (x-axis) versus L-NAME infusion (y-axis). L-NAME caused a significant decrease in arterial diameter (−10.6±4.6, LME p=7.6×10^−6^ Bonferroni corrected, n=6 mice, 36 vessels). **I.** Population locomotion-triggered average after vehicle (black) and L-NAME (red) infusions (n=6 mice, 26 arterioles in FL/HL representation). For both cases, the diameters are normalized by the average basal diameter of the vessel after vehicle infusion. **J.** LFP power spectra during stationary (basal) and locomotion periods after L-arginine infusion, normalized to vehicle infusion in the same mouse. L-arginine slightly reduces neural activity in the gamma band (basal, −25.7±3.1%, paired t-test p=0.13; locomotion, −3.4±6.9%, paired t-test p=0.80, n=3). **K.** Plot of basal arteriole diameter after vehicle infusion (x-axis) versus L-arginine infusion (y-axis). There was no significant change in the arterial diameter (7.1±4.9%, LME p=0.43 Bonferroni corrected, n=5 mice, 24 vessels). **L.** Population locomotion-triggered averages after vehicle (black) and L-arginine (red) infusions (n=5 mice, 18 arterioles in FL/HL representation). For both cases, the diameters were normalized by the average basal diameter of the vessel after vehicle infusion. **M.** Representative image of cortex taken of AAV-hSyn-DIO-hM3D(Gq)-mCherry in nNOS-cre mice, where DREADDs expressed in nNOS^+^ cells. Left image is wide field image of nNOS-mCherry DREADDs virus (red) in cortex with DAPI (blue) staining (scale bar 100μm). Right image is zoomed image of nNOS-mCherry DREADDs virus (red) in cortex with DAPI (blue) and GAD65 (green) staining (scale bar 30μm). **N.** Comparison of the percent change in basal arteriole diameter after treatment with L-NAME (n=6 mice, 36 vessels) and nNOS-G(i) DREADDs (n=9, 49 vessels) relative to vehicle controls (aCSF infusions and DMSO injection, respectively). The percent changes to the basal vessel diameter are not significantly different (LME p=0.57), showing the NO regulation of basal vessel diameter is primarily produced by nNOS in neurons, not eNOS from endothelial cells. **O.** Comparison of the percent change in basal arteriole diameter after treatment with L-arginine (n=5 mice, 24 vessels) and nNOS-G(q) DREADDs (n=6, 32 vessels). The percent changes to the basal vessel diameter were significantly different (LME p=0.02).

We then tested whether the effects of nNOS-expressing neuron activity on arteriole diameter was mediated by NO, as these neurons also release vasoactive peptides^17^. Previous studies with NOS inhibitors have shown conflicting effects^47, 48^. Discrepancy in the literature likely results from differences in experimental methodology: topical application does not reach lower cortical layers^35^ where most of the nNOS expressing neurons reside, and systemic administration impacts the cardiovascular system. To avoid these experimental confounds, we infused the water-soluble NO synthase inhibitor, Nω-nitro-L-arginine methyl ester (L-NAME), or the precursor for NO, L-arginine, through an implanted cannula to decrease or increase NO-mediated signaling, respectively. Note that previous work has shown that cortical neurons express eNOS^49–51^, endothelial cells on cerebral arterioles express nNOS^37^, and that there are no truly specific pharmacological inhibitors of nNOS^52–55^. Locomotion-evoked dilations are unlikely to be mediated by eNOS expressed in endothelial cells because shear-mediated increases in eNOS activity in response to flow-evoked changes in diameter evolve over minutes, not seconds^56^, too slow to account for the locomotion-evoked dilations. Blocking the production of NO caused no significant change in the basal gamma-band power (+8.24±2.8%, paired t-test p=0.39, Fig. 3G), but a strong decrease in the basal vessel diameter and locomotion-evoked dilation (−10.6±4.6%, LME p=7.6×10^−6^, Fig. 3H and −9.6±1.9%, paired t-test p=5.2×10^−4^, Fig. 3I). Infusion of L-arginine, did not significantly affect gamma-band activity (− 25.7±3.1%, paired t-test p=0.13, Fig. 3J), and caused a non-significant increase in basal arterial diameter (+7.1±4.9%, LME p=0.43, Fig. 3K). Comparisons of the average constriction induced by L-NAME to the constriction induced by hM4D-G(i) DREADDs in nNOS-expressing neurons and showed no significant difference (Fig. 3N, t-test p=0.57), indicating that the NO regulating basal vessel diameter is primarily produced by nNOS in neurons, not eNOS from endothelial cells.

There are two varieties of nNOS–expressing neurons in the cortex, Type 1 and Type 2^57^. Type 2 nNOS neurons are a heterogeneous group of interneurons that are more numerous and located throughout the cortex. In contrast, Type 1 nNOS neurons are sparse, are located in deeper layers (similar to our viral expression, Fig. 3M), and express nNOS at higher levels. While Type 1 nNOS neurons receive many different neuromodulatory inputs^58, 59^, nearly all of them express the Substance P (SP) receptor (NK1R), and they are the only cells in the cortex that do so^60–62^. Type 1 nNOS neurons also have extensive, long-range projecting axonal arbors that well-position them to influence the vasculature. Substance P causes prolonged depolarization and spiking in Type 1 nNOS neurons^62^. As there are no NK1 receptors in vascular cells^37^, we can pharmacologically manipulate the activity of Type 1 nNOS neurons independently of other neurons by infusing SP, or an antagonist for the SP receptor, CP-99994. Infusions of CP-99994 caused non-significant decreases in gamma-band power (−5.28±5.11%, paired t-test p=0.5 Fig. 4A) and a non-significant decrease in basal arteriole diameter (−8.0±7.2%, LME p=1 Fig. 4B). Infusions of SP caused non-significant decreases in gamma-band power (−24.42±3.8%, paired t-test p=0.19 Fig. 4D), but a substantial increase in basal diameter (+13.4±4.4 %, LME p=4.2×10^−3^ Fig. 4E). Co-infusion of SP with muscimol produced no dilation (Fig. S9), consistent with the action of SP being mediated through local neural excitation. These results show that NO released from a small subset of neurons, whose activity is not reported in measures of population activity, controls both the basal diameter of arterioles, and mediates a substantial portion of the hyperemic response.

**Figure 4.**
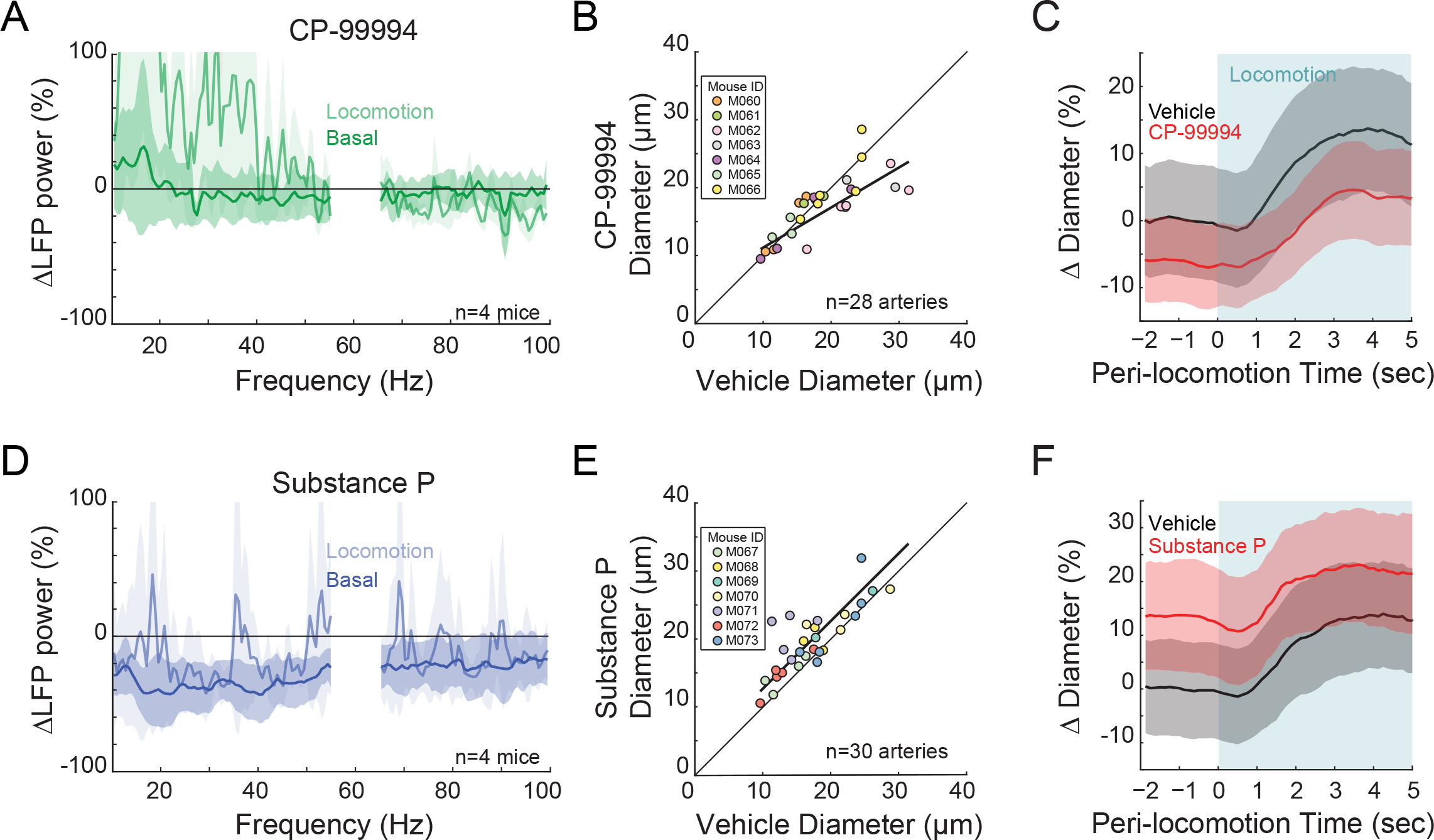
Type I nNOS expressing neurons control basal arteriole diameter. Data in **A-C** are from mice after CP-99994 infusions, which will block the excitatory Substance P input onto Type1 nNOS neurons, data in **D-E** is from mice after Substance P infusions, which will excite Type1 nNOS neurons. **A.** LFP power spectra during stationary (basal) and locomotion periods after CP-99994 infusion, normalized to vehicle infusion in the same mouse. CP-99994 non-significantly reduced neural activity in the gamma band (basal, −5.28±5.11%, paired t-test p=0.5; locomotion, −6.8±52.3%, paired t-test p=0.69, n=4). Shading denotes mean ± standard deviation. **B.** Plot of basal arteriole diameter after vehicle (x-axis) versus CP-99994 infusion (y-axis). CP-99994 caused a non-significant decrease in the basal arterial diameter (−8.0±7.2%, LME p=1, n=7 mice, 28 vessels). **C.** Population locomotion-triggered averages after vehicle (black) and CP-99994 (red) infusions (n=7 mice, 14 arterioles in FL/HL representation). For both cases, the diameters were normalized by the average basal diameter of the vessel after vehicle infusion. **D.** LFP power spectra during stationary (basal) and locomotion periods after Substance P infusion, normalized to vehicle infusion in the same mouse. Substance P causes a non-significant reduction of neural activity in the gamma band (basal, −24.42±3.8 %, paired t-test p=0.19; locomotion, −12.7±4.2 %, paired t-test p=0.67, n=4). **E.** Plot of basal arteriole diameter after vehicle infusion (x-axis) versus Substance P infusion (y-axis). Substance P caused a significant increase in the basal arterial diameter (+13.4±4.4 %, LME p=4.2×10^−3^ Bonferroni corrected, n=7 mice, 30 vessels). **F.** Population locomotion-triggered averages after vehicle (black) and Substance P (red) infusions (n=7 mice, 16 arterioles in FL/HL representation). For both cases, the diameters were normalized by the average basal diameter of the vessel after vehicle infusion.

### Relationship between neural activity and arterial diameter changes in the basal and evoked conditions

To better understand the overall relationship between neural activity and arterial diameter, we plotted all the impacts of all of our perturbations (Fig. 5A). If arterial diameter tracks neural activity, then we would expect all the points to lie in a straight line or along a curve. However, we see no one-to-one relationship between basal neural activity and basal arterial diameter. A similar lack of one-to-one relationship between neural activity and dilation is also seen during locomotion (Fig. 5B). Perturbations of neural activity or NO production can drive large changes in arterial diameter that are incommensurate, or even opposing what one would expect from the neural activity. With the exception of muscimol, locomotion drives a similar increase in neural activity across all perturbations, though the amount of vasodilation is cut in half when the production of NO is blocked at the cellular (G(i) DREADDs, CP99994) or biochemical (L-NAME) level, an indication that NO signaling from neurons drives approximately half of the functional hyperemic response. Importantly, the manipulations of NO diameter up (G(q) DREADDs, L-arginine and Substance P) or down (G(i) DREADDs, L-NAME, CP-99994) using multiple orthogonal approaches results in very similar changes in neural activity and arterial diameter, suggesting that a small nexus (an ‘oligarchy’) of NO-producing Substance P-responsive neurons are the primary controllers of the cerebral vasculature, not the average activity of all neurons.

**Figure 5.**
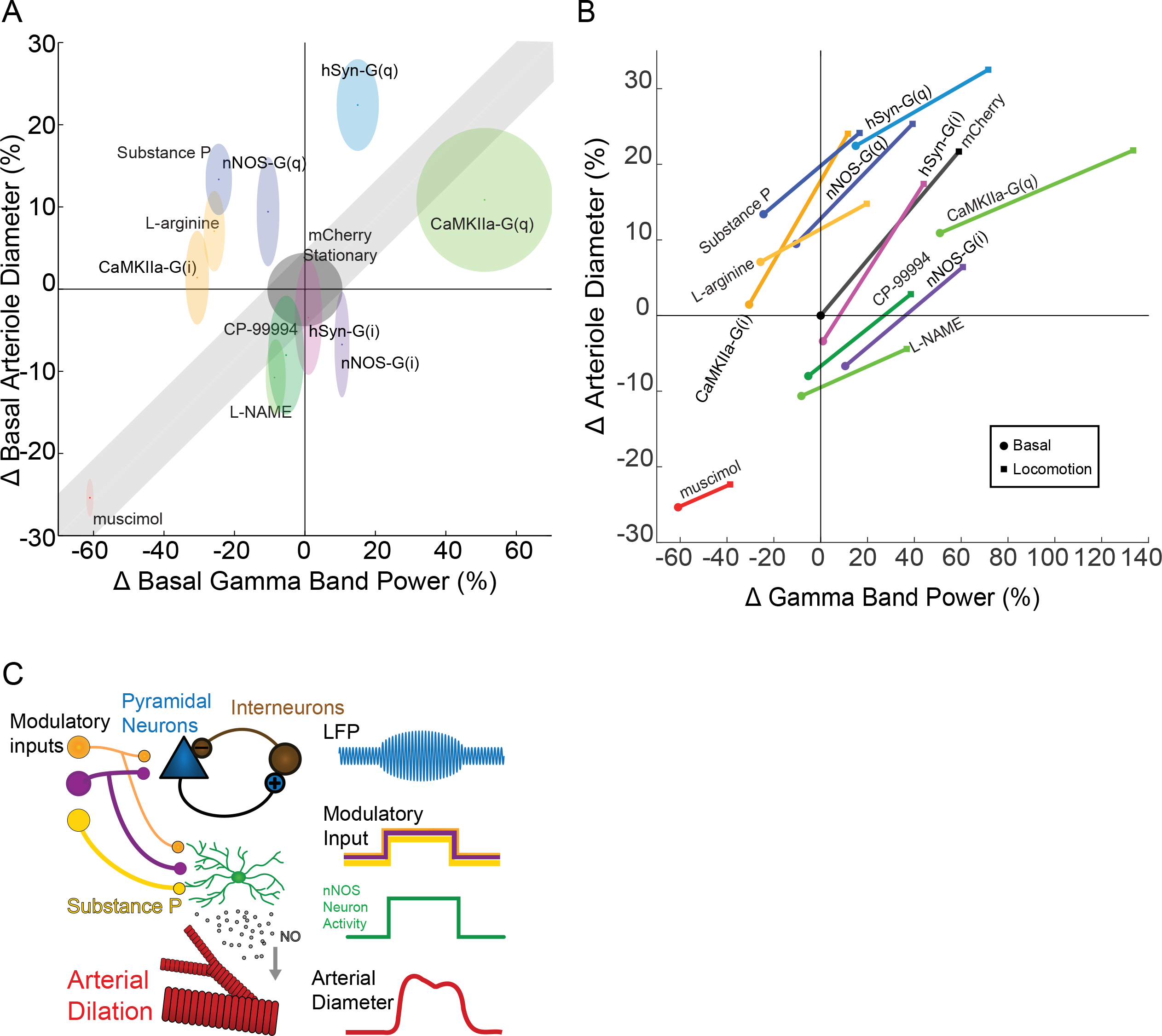
Lack of one-to-one relationship between neural activity and arterial dilation. **A.** Summary figure outlining the change in basal gamma band power vs change in basal arteriole diameter under each condition outlined in this paper. Shading surrounding each point shows standard deviation from mean. The unity line represents where the points would lie if there was a linear relationship between neural activity and vasculature. For chemogenetic manipulations, neural activity and basal arterial dilation have been shifted so that the reporter virus expressing animals (mCherry) are centered at the origin. **B.** Summary figure plotting the change in gamma band power vs in arteriole diameter during locomotion for the various manipulations performed. Note the lack of a one-to-one relationship between neural activity change and arterial dilation. **C.** Schematic showing neural circuit which can account for the data. Pyramidal and nNOS type 1 neurons are both driven by common modulatory drive, resulting in a correlation between gamma-band power increases and vasodilation during normal behavior.

## Discussion

Our results show that both basal and sensory-evoked changes in arterial diameter are controlled by local neural activity. Surprisingly, large alterations of the activity of pyramidal neurons had little effect on either basal or evoked arterial diameter, even though the activity of these neurons dominates extracellular recordings of electrical signals and are the major energy consumers in cortex^43^. Rather, cortical hemodynamic signals are controlled primarily by NO produced by the activity of nNOS-positive neurons. Our experiments with NOS-inhibitor infusion and suppression of nNOS-expressing neural activity with DREADDs showed that approximately half of the evoked arterial dilatory response, and half of the baseline artery diameter change seen with muscimol infusion, were caused by neurally-generated NO. Other neurovascular coupling mechanisms such as voltage-gated potassium channels^44^ and signaling from astrocytes^63^ likely underlie the remaining basal and evoked diameter changes.

Hemodynamic signals have generally been interpreted as reporters of bulk neural activity (but see^5^), with increases in neural activity thought to be directly correlated with increases in vessel dilation and blood flow. Our results support an alternative model, where basal and evoked blood flow is controlled by a selected group of neurons independent of the bulk population activity. Rather than being a ‘democracy’, where the activity of all neurons contributes to increases in blood flow, vascular control in the cortex is an ‘oligarchy’, where a small group of neurons controls the flow largely independent of the activity of their neighbors. This is apparent when we look at the relationship between gamma-band power and basal (Fig. 5A) or evoked arteriole diameter (Fig. 5B). We see no one-to-one relationship between neural activity and the hemodynamic response, meaning local neural activity cannot always be decoded from hemodynamic signals. We note that this problem is exacerbated when using BOLD imaging, as changes in baseline blood flow can change the magnitude or even the sign of the BOLD response without changes in the underlying neural activity^23, 24^.

Our data point to a way of resolving several troubling issues in the field of neurovascular coupling. The mismatch between neural activity and hemodynamic signals, seen in many studies^9, 64^, can be resolved if the hemodynamic signals are not controlled by the overall local neural activity, but by a small group of neurons primarily driven by modulatory signals that are engaged when a sensory stimulus is presented or by attention modulation of neural activity^11^ (Fig. 5C). The lack of effect of changes in pyramidal neuron activity on arterial diameters suggests that these type 1 nNOS neurons receive minimal local recurrent excitatory input. At least in the primary sensory cortex, during natural behaviors and passive sensory stimulation, the activity of the nNOS-positive neurons and pyramidal neurons are correlated, generating the widely observed linkage between vasodilation and gamma band power^6, 7, 65^ (Fig. 1), though because of the small size and number of Type 1 nNOS neurons (<1% of all neurons) and their closed electrical fields, their activity will not be detectable in the local field potential^66^ or in single unit recordings. Stimuli or modulatory conditions that cause the activity of nNOS-positive neurons to become uncorrelated from the activity pyramidal neurons will drive neurovascular decoupling, as has been observed in many studies^9, 10, 33^. Type 1 nNOS neurons are hubs for modulation, as they receive orexinergic and substance P input, as well as cholinergic input from the basal forebrain^58, 67–69^. Stimulation of the basal forebrain is known to greatly increase basal cortical blood flow independent of metabolic changes^70, 71^, and the projections of basal forebrain neurons are topographically mapped^72, 73^, consistent with the spatially localized hemodynamic responses. Increases in the activity of cholinergic neurons in the basal forebrain and other modulatory regions occur during sensory stimulation^74^ and voluntary movement^75^, consistent with the vasodilation accompanying these behaviors^7, 33^. These neurons may also play a role in the large increases in cerebral blood flow seen during REM sleep^76, 77^.

As our works shows that cerebral blood flow is controlled by a relatively small subset of neurons, and not by the average population activity, any damage or dysfunction to this select group of neurons could result in decreased blood flow, regardless of the metabolic need. Importantly, changes in activity patterns of these neurons will alter basal diameter of arteries, which is known to affect the amplitude of fMRI signals^24^. There is a decline in the number of interneurons, particularly nNOS-expressing interneurons, with aging^78, 79^, and the removal of their tonic vasodilatory signal could cause decreases in cerebral blood flow that are thought to contribute to dementia^25^. Indeed, alterations of nNOS and TAC1 (encoding the precursor to Substance P) expression have been implicated in Alzheimer’s disease^80, 81^. Protecting these blood-flow controlling neurons from insults and damage could be a promising strategy for preventing neurovascular disorders.

## Methods

### Animal Procedures

All procedures were performed in accordance with protocols approved by the Institutional Animal Care and Use Committee (IACUC) of Pennsylvania State University. Data were acquired from 165 mice (95 mice for 2PLSM imaging, 61 mice for electrophysiology, and 20 mice for immunohistochemistry - see Tables 1 and 2) C57-BL6J mice (Jackson Laboratory) and cre-nNOS mice (B6.129-*Nos1^tm1(cre)Mgmj^*/J) between 3-8 months of age. Mice were given food and water ad libitum and maintained on 12-hour light/dark cycles in isolated cages during the period of experiments. All *in vivo* experiments were performed in the morning.

**Table 1.** AAV-injected mice

| Table 1: AAV-injected mice | # mice (vessels) for 2-photon imaging | # mice for chronic electrophysiology | Immunohistochemistry |
| --- | --- | --- | --- |
| C57-BL6J: AAV5-CMV-TurboRFP-WPRE-rBG (UNC Vector Core) | 12 mice (56 vessels) | 4 mice | 3 mice |
| C57-BL6J: AAV8-hSYN-HA-hM3D(Gq)-mCherry (Addgene #50474-AAV8) | 7 mice (37 vessels) | 3 mice | 3 mice |
| C57-BL6J: AAV5-hSYN-HA-hM4D(Gi)-mCherry (UNC Vector Core) | 7 mice (45 vessels) | 3 mice | 2 mice |
| C57-BL6J: AAV5-CaMKIIa-hM3D(Gq)-mCherry (Addgene # 50476-AAV5) | 6 mice (41 vessels) | 4 mice | 2 mice |
| C57-BL6J: AAV5-CaMKIIa-hM4D(Gi)-mCherry (UNC Vector Core) | 7 mice (38 vessels) | 5 mice | 2 mice |
| B6.129-*Nos1<sup>tm1(cre)Mgmj</sup>/J*: AAV8-hSyn-DIO-hM3D(Gq)-mCherry in nNOS-cre mice (Addgene #44361-AAV8) | 6 mice (32 vessels) | 4 mice | 2 mice |
| B6.129-*Nos1<sup>tm1(cre)Mgmj</sup>/J*: AAV8-hSyn-DIO-hM4D(Gi)-mCherry in nNOS-cre mice (Addgene #44362-AAV8) | 9 mice (49 vessels) | 5 mice | 2 mice |

**Table 2.** Infused mice

| Table 2: Infused mice | # mice (vessels) for 2-photon imaging | # mice for chronic electrophysiology |
| --- | --- | --- |
| C57-BL6J: Muscimol (10mM) | 6 mice (26 vessels) | 3 mice |
| C57-BL6J: L-Arginine (10μM) | 5 mice (24 vessels) | 3 mice |
| C57-BL6J: L-NAME (1mM) | 6 mice (36 vessels) | 5 mice |
| C57-BL6J: Substance P (1μM) | 7 mice (30 vessels) | 4 mice |
| C57-BL6J: CP-99994 (8mM) | 7 mice (28 vessels) | 4 mice |
| C57-BL6J: ML-133 (10μM) | 5 mice (36 vessels) | 4 mice |
| C57-BL6J: BaCl<sub>2</sub> (100μM) | 5 mice (24 vessels) | 3 mice |

#### Virus Injections

All viruses were obtained from UNC Vector Core or Addgene (see Table 1). For viral injections, mice were anaesthetized with isoflurane (5% induction, 2% maintenance) and placed in a stereotaxic rig. A small craniotomy was made over the FL/HL representation of the somatosensory cortex (0.75 mm caudal, 2.5 mm lateral from bregma). Using a pulled glass micropipette (30-50μm tip diameter) and infusion pump (Harvard Apparatus, Holliston, MA), we microinjected 500 nanoliters (at a rate 100nl/min) of the adeno-associated viral (AAV) vectors (see Table 1) ~300μm below the cortex. After the viral injections were completed, the glass micropipette was kept in place for 10 minutes before removal. The skin was sutured, and mice were returned to their home cage. Four weeks after AAV injections, we performed surgeries for electrode or window implantation, or perfusion for immunohistochemistry.

#### Window, electrode, and cannula implantation procedures

Mice were anesthetized with isoflurane and the scalp was resected. A custom-made titanium metal bar was fixed to the skull with cyanoacrylate glue (Vibra-Tite, 32402) just posterior to the lambda cranial suture. A 4-8 mm^2^ area polished and reinforced thinned-skull window was created over the forepaw/hindpaw representation of somatosensory cortex on the right hemisphere, as described previously^7, 27^. The thinned area was polished with size 3F grit (Covington Engineering, Step Three 3F-400) and reinforced with a fitted #0 glass coverslip (Electrode Microscopy Sciences, #72198). Self-tapping screws (#000, 3/32”, JI-Morris, Southbridge, MA) were placed into the contralateral parietal and ipsilateral frontal bone to stabilize the skull. The titanium bar and screws were secured with black dental acrylic resin. The animals were allowed 2-3 days to recover before habituation.

For electrode or cannula implantation, small (<0.5mm diameter) craniotomies were made to insert the stereotrode or cannula (Plastics One, C315DCS, C315GS-4) into the upper layers of cortex near the FL/HL representation of cortex. The cannula was attached to the skull and headbar with cyanoacrylate glue and dental acrylic. Implanted cannulas do not alter the hemodynamic response^7^.

#### Habituation

Mice were habituated to head-fixation on the spherical treadmill (60 mm diameter) over three days before imaging. Mice were habituated for 15 minutes of acclimation on the first day. On successive days, the time was increased to up to two hours. During habituation the mice were monitored for any signs of distress during habituation. Optical imaging data were taken over 4 imaging sessions (two sessions under each condition for the 2PLSM).

#### Intraperitoneal injections and cortical infusions

CNO (2.5 mg/kg in 2% DMSO in saline) or vehicle were injected intraperitoneal 20 minutes before imaging began. Electrophysiological experiments (data not shown) showed that the neural effects of DREADDs onset within 20 minutes of injection. CNO and DMSO were each injected in a counterbalanced order.

For the intracortical infusions, mice were first head-fixed on the spherical treadmill. The dummy cannula was then slowly removed and replaced with an infusion cannula (Plastics One, C315IS-4). The interface between the infusion cannula and the guide cannula was sealed with Kwik-Cast (World Precision Instruments). All infusions were done at a rate of 25 nL/min for a total volume of 500 nL. Animals were kept in place for 20 min after the end of the infusion and then moved to the imaging apparatus. The infusion cannulas were kept in place for the duration of the imaging session. Pharmacological treatments and aCSF were each infused in a counterbalanced order.

### Physiological measurements

#### Two-photon microscopy

Prior to imaging, mice (n=95 mice) were briefly anesthetized with isoflurane, and retro-orbitally injected with 50 μL 5% (weight/volume) fluorescein conjugated dextran (FITC-dextran 70 kDa; Sigma-Aldrich)^27, 40^ then head-fixed upon a spherical treadmill. The treadmill was coated with anti-slip tape and attached to an optical rotary encoder (US Digital, E7PD-720-118) to monitor rotational velocity. Velocity changes were used to distinguish between periods of rest and locomotion. Imaging was done on a Sutter Movable Objective Microscope with a 16x 0.8NA objective (Nikon). A MaiTai HP laser tuned to 800 nm was used to excite the FITC. The power exiting the objective was 10-15 mW for imaging surface arteries. For vessel diameter measurements, movies of individual arteries were taken at a nominal frame rate of 8 Hz for 5 minutes. For RBC velocity measurements line-scans were made along the long axis of the capillary lumen (Fig. S4A)^28^. The same vessel segments were imaged after both the treatment (intraperitoneal CNO injection or local drug infusion) and vehicle controls (intraperitoneal vehicle injection or intracranial vehicle infusion). Treatments and vehicle condition were imaged in a counterbalanced order on separate days. Imaged arteries were chosen in the first session of imaging. Arteries were visually identified by their more rapid blood flow, rapid temporal dynamics of their response to locomotion, and vasomotion^1, 7, 27^. Examples of individual vessels for each condition with and without manipulations are seen in Fig. S10. Bright field and fluorescent images of the PoRTs window were taken before imaging on the 2-photon for each mouse. For imaging experiments with DREADDs manipulations the imaged vessels were chosen if they were within the fluorescent regions of the PoRTs window (Fig. 2A). For infusion imaging experiments, the imaged vessels were chosen if they were within 1.5mm of the cannula. For repeated imaging of the same vessel, three-dimensional image stacks of the regions were made, and the position of nearby vessels were used to return to the same imaging plane on subsequent imaging sessions. Only vessels anatomically identified to be in the FL/HL representation were used in locomotion-triggered averages.

#### Image processing and data analysis

Data analysis was performed in MATLAB (MathWorks). 2PLSM images were aligned in the x–y plane using a rigid-body registration algorithm^1, 33, 82^. Each movie was manually reviewed to ensure that z-axis motion during locomotion was minimal. A rectangular box was manually drawn around a segment of the vessel. For surface arteries, the intensity of a short segment (1-3 micrometers in length) was averaged along the long axis of the vessel, and the diameter was calculated from the full-width at half-maximum^1^. For penetrating arteries, the Thresholding in Radon Space (TiRS) method^83^ was used as it effectively performs a full-width at half maximum measurement along every angle of the penetrating artery (https://github.com/DrewLab/Thresholding_in_Radon_Space). For diameters of penetrating vessels, using the TiRS method is important as penetrating vessels are not circular and the cross sectional shape will change during dilation, making single-axis diameter measurements inaccurate^83–85^. A square region of interest (ROI) enclosing the penetrating arteriole of interest was manually drawn. The images were transformed into Radon space, thresholded, and then transformed back to image space, where the vessel cross-sectional area was quantified after a second thresholding. In order to facilitate comparison with pial vessels, diameters (D) of penetrating vessels were taken to be: 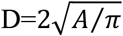, where A was the cross-sectional area calculated using the TiRS method. Frames in which the temporal derivative of diameter changed by >16 μm/s were tagged as motion artifacts and replaced with the linear interpolation between the proceeding and subsequent points. The diameter was filtered with a five-point median filter (MATLAB function: medfilt1; filter order = 5). Red blood cell velocity in capillaries was calculated using the Radon transform^40^ (https://github.com/DrewLab/MCS_Linescan). Example images of individual vessels were 3D median filtered in ImageJ (ImageJ: 3D Median, radius=2) and then the max intensity projection was taken for 40 frames with or without locomotion.

To detect locomotion events, the signal from the rotational encoder on the spherical treadmill was low-pass filtered (10 Hz, fifth-order Butterworth), and then the absolute value of the acceleration was binarized with a 10^−5^ cm/s^2^ threshold^27, 33^. Stationary periods were defined as times when the mouse was still, with a 2s buffer after the end of any preceding locomotion event and a 1s buffer before the start of the next locomotion event. Onset time was calculated with a linear regression between 20% and 80% to the peak dilation^86^. The change in basal diameter in each treatment condition was normalized by the basal diameter of the same vessel in the vehicle control condition.

For the vessels that were histologically identified as being within the FL/HL representation, locomotion-triggered averages (LTA) were calculated by taking the average of any locomotion event >5 seconds in duration with at least 2 seconds of no locomotion before the event. These averages were normalized by the average basal diameter of the same vessel under vehicle control conditions.

#### Electrophysiology

Neural activity was recorded as differential potentials between the two leads of Teflon-insulated tungsten micro-wires (A-M Systems, #795500)^7, 8, 33^. Differential recordings between two closely spaced electrodes avoid the volume conduction of remote signals^87^. Stereotrode micro-wires, with an inter-electrode spacing of ~100μm, were threaded through polyimide tubing (A-M Systems, #822200). Electrode impedances were between 70-120 kΩ at 1 kHz. The electrodes were implanted in the upper layers of cortex (~400 μm depth). The acquired signals were amplified (World Precision Instruments, DAM80), band-pass filtered between 0.1 and 10 kHz (Brownlee Precision, Model 440) during acquisition, then digitized at 20 kHz.

The local field potential (LFP) was calculated by digitally band-pass filtering the raw neural signal between 10-100 Hz (MATLAB function: butter, filtfilt; filter order = 4) and using a notch filter to remove 60 Hz noise (MATLAB function: iirnotch). The basal power was computed by averaging the power during periods of recording with no locomotion. Normalized power for each recording session was calculated by normalizing by the basal power after vehicle injection/infusion in the same mouse. Gamma-band power was quantified by averaging the 40-100 Hz band of the LFP after using a notch filter to remove 60 Hz noise.

We quantified the multi-unit average (MUA) by taking the power of the signal within the 300-3000 Hz band^6^. Power was calculated after digitally band-pass filtering the raw neural signal (MATLAB function: butter, filtfilt; filter order = 4). The envelope of this signal was quantified using the Hilbert transform.

### Histology

At the end of the experiment, mice were deeply anesthetized with 5% isoflurane, and then transcardially perfused with heparinized saline followed by 4% paraformaldehyde. The brain was removed from the skull and sunk in 30% sucrose/4% PFA. For cytochrome oxidase staining, the cortex was then flattened and 60μm tangential sections were cut on a freezing microtome. The cytochrome oxidase in the sections was stained, and the location of the FL/HL representation in somatosensory cortex was reconstructed relative to the vasculature visible through the cranial window^30^.

For immunohistochemistry labeling, the brains were immersed in a 4%PFA/30% sucrose solution for 1 day. Tissues were sectioned coronally to a 90μm thickness and then washed in PBS twice. For heat induced antigen retrieval, the slices were boiled in 10mM sodium citrate for 10 minutes. The sections were then incubated with primary antibodies on slides at a 1:200 dilution for two days at 4°C (Mouse monoclonal GAD-65 Antibody (A-3): sc-377145 Santa Cruz Biotechnology, Mouse monoclonal CaMKII Antibody (G-1): sc-5306 Santa Cruz Biotechnology, Rabbit polyclonal to GFAP: ab7260 abcam). The brain sections were washed twice and incubated for one hour with secondary antibodies (Abcam Alexa Fluor^®^ 488 Goat Anti-Mouse IgG H& L and Goat Anti-Rabbit IgG H& L). The brain sections were washed twice, and coverslipped with DAPI fluoroshield mounting media (Abcam ab104139). The sections were imaged on an Olympus FV10i Confocal.

### Statistical analysis

Statistical analysis was performed using MATLAB (R2018, MathWorks). All summary data were reported as mean ± standard deviation. For comparisons of vehicle vs. treatment conditions, where multiple measurements of different vessel diameters were taken from the same mouse, we used the linear mixed effects (LME) model (MATLAB function: fitlme). For each manipulation (e.g. CNO vs. vehicle in DREADD-expressing mice) we fit a linear mixed effects model given by:

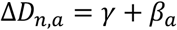

where Δ*D*_*n, a*_ is the diameter differences between the vehicle and manipulation condition, *a* is the identifier of the animal group, and *n* is the identifier of the vessel. The *γ*, term is the (fixed) effect of the manipulation, and *β*_*a*_ is the within animal (random) effect that accounts for the within animal correlations^29^. To determine if the conditions showed a significant change in the vessel diameter, we calculated if the *γ* term differed from 0, after a Bonferroni correction for the number of groups of DREADD-expressing mice (7 groups) or the number of infusions (7 groups). A difference was determined to be statistically significant if p<0.05 after Bonferroni correction. For plotting the relationship of the basal diameter in the control condition versus the manipulation in the figures, we fit a linear mixed effects model given by:

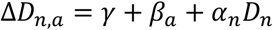

where *D*_*n*_ is the diameter of vessel *n* in the control condition, and *α*_*n*_ (is a vector of (fixed) coefficients.

For comparisons of vehicle vs. treatment conditions of the gamma-band (averaged power from 40-100Hz after removing 60 Hz noise), we used the paired t-test (MATLAB function: ttest).

---

**Supplementary Figure 1.**
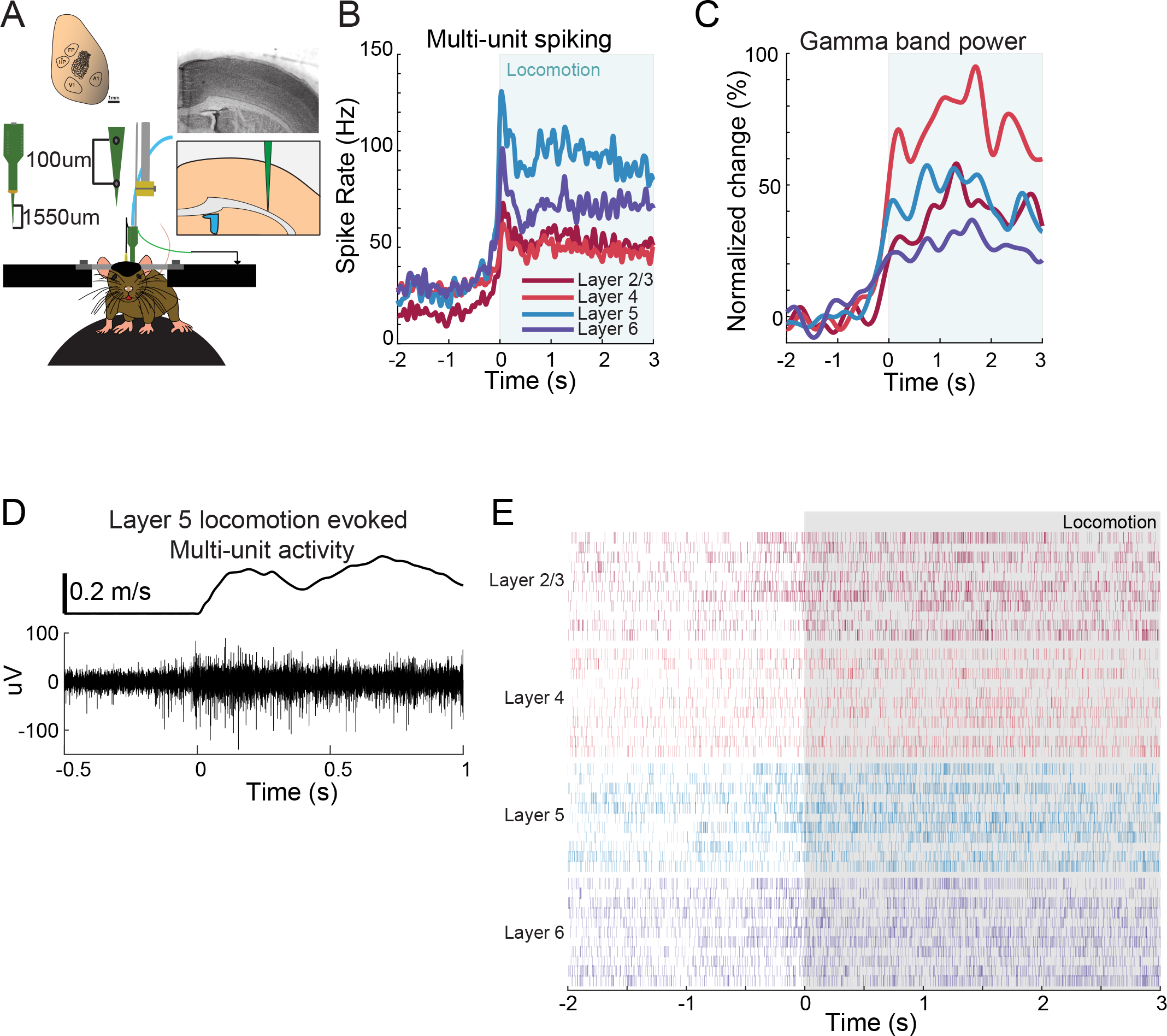
Example neural responses in forelimb/hindlimb representation of somatosensory cortex during voluntary locomotion. **A.** Schematic showing experimental setup of electrophysiological measurements taken with multi-electrode array. **B.** Average locomotion-evoked multi-unit spiking from a single site in the FL/HL representation in S1. **C.** Average locomotion-evoked gamma-band power change from a single site in the FL/HL representation in S1. **D.** Example of multi-unit activity recorded in layer 5 during locomotion. **E.** Locomotion-triggered spike rasters for several locomotion events.

**Supplementary Figure 2.**
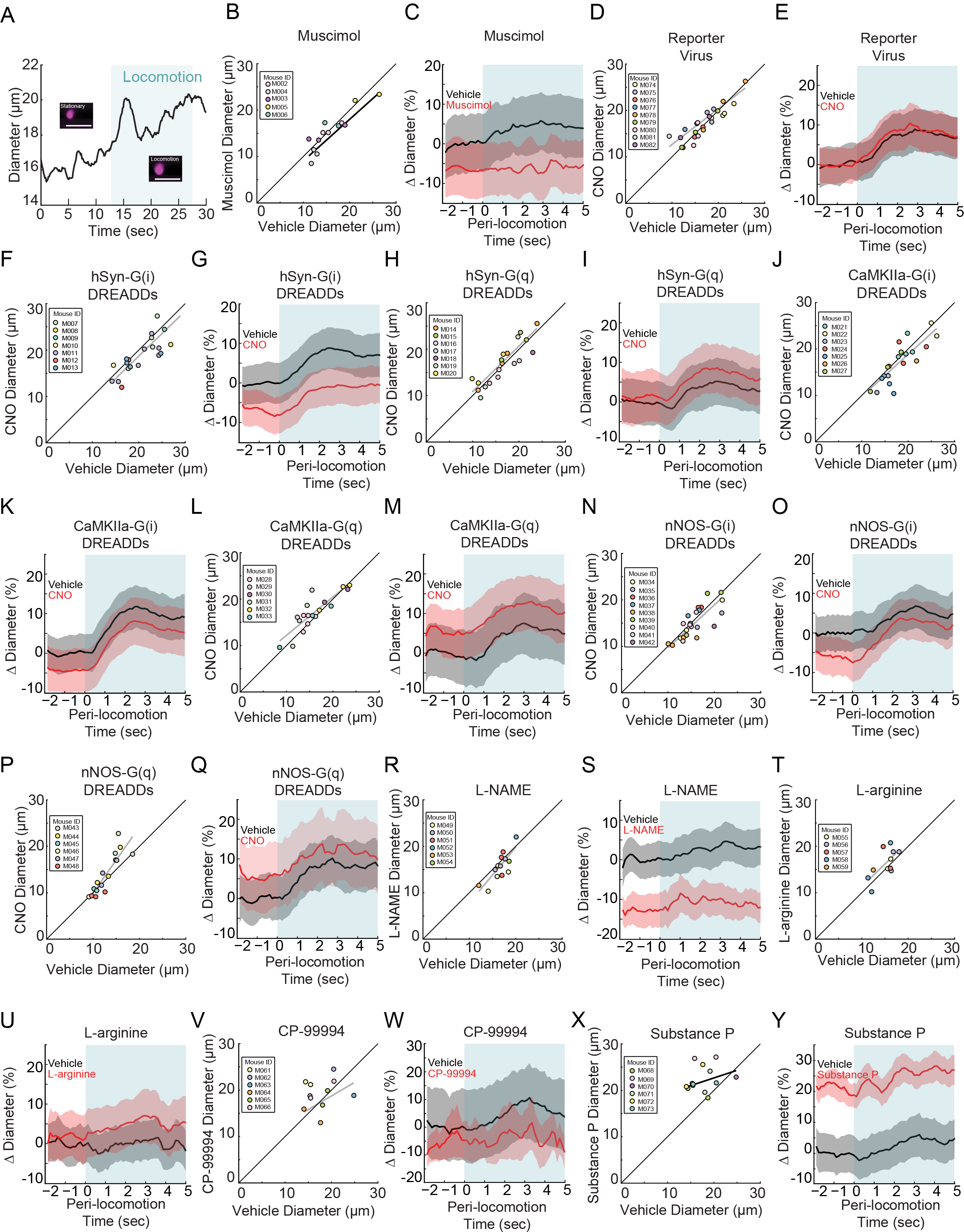
Locomotion-triggered dilations of penetrating arterioles. **A.** Example of locomotion-induced dilation in a penetrating arteriole. Inset shows the vessel during a stationary period and during locomotion. Blue shading denotes periods of locomotion. Data in **B** and **C** are from penetrating arteries in mice infused with muscimol and aCSF. **B.** Basal diameter of penetrating arteries after vehicle infusion (x-axis) versus muscimol infusion (y-axis). There was not a significant decrease in basal diameter in the muscimol infused condition (−7.8±1.4 %, LME p=1 Bonferroni corrected, n=5 mice, 13 vessels). **C.** Population locomotion-triggered averages after vehicle (black) and muscimol (red) infusions (n=5 mice, 13 arterioles in FL/HL representation). For both cases, the diameters were normalized by the average basal diameter of the vessel after vehicle infusion. Shading denotes mean ± standard deviation. Data in **D** and **E** are from penetrating arteries in mice infected with AAV-CMV-TurboRFP-WPRE-rBG (reporter virus), and IP injected with CNO or vehicle control. **D.** Basal arteriole diameter of penetrating arteries after vehicle injection (x-axis) versus CNO injection (y-axis) was not significantly different (−0.3±14.2 %, LME p=1 Bonferroni corrected, n=5 mice, 13 vessels). **E.** Population locomotion-triggered averages in response to 5 seconds of locomotion after vehicle (black) and CNO (red) injection (n=9 mice, 13 arterioles in FL/HL representation). Data in **F** and **G** are from penetrating arteries in mice infected with AAV-hSYN-HA-hM4D(Gi)-mCherry, data in **H** and **I** is from penetrating arteries in mice infected with AAV-hSYN-HA-hM3D(Gq)-mCherry (pan-neuronal expression) **F.** Basal diameter of penetrating arteries after vehicle injection (x-axis) versus CNO injection (y-axis) was not significantly decreased by CNO (−6.7±12.0 %, LME p=0.19 Bonferroni corrected, n=7 mice, 21 vessels). **G.** Population locomotion-triggered averages after vehicle (black) and CNO (red) injection (n=7 mice, 21 arterioles in FL/HL representation). **H.** Basal diameter of penetrating arteries after vehicle injection (x-axis) versus CNO injection (y-axis). There was no significant difference between the conditions (+2.3±13.9 %, LME p=1 Bonferroni corrected, n=7 mice, 19 vessels). **I.** Population locomotion-triggered averages after vehicle (black) and CNO (red) injection (n=7 mice, 19 arterioles in FL/HL representation). Data in **J** and **K** are from penetrating arteries in mice infected with AAV-CaMKIIa-hM4D(Gi)-mCherry, data in **L** and **M** is from penetrating arteries in mice infected with AAV-CaMKIIa-hM4D(Gq)-mCherry (expression in pyramidal neurons) **J.** Basal arteriole diameter of penetrating arteries after vehicle injection (x-axis) versus CNO injection (y-axis) showed no significant difference (−4.1±13.5 %, LME p=0.42 Bonferroni corrected, n=7 mice, 20 vessels). **K.** Population locomotion-triggered averages after vehicle (black) and CNO (red) injection (n=7 mice, 20 arterioles in FL/HL representation). **L.** Basal arteriole diameter of penetrating arteries after vehicle injection (x-axis) versus CNO injection (y-axis) showing no significant difference (+8.5±14.6 %, LME p=0.30 Bonferroni corrected, n=7 mice, 18 vessels). **M.** Population locomotion-triggered averages after vehicle (black) and CNO (red) injection (n=7 mice, 18 arterioles in FL/HL representation). Data in **N** and **O** are from penetrating arteries in nNOS-cre mice injected with AAV-hSyn-DIO-hM4D(Gi)-mCherry, data in **P** and **Q** is from penetrating arteries in nNOS-cre mice injected with AAV-hSyn-DIO-hM4D(Gq)-mCherry (nNOS+ expression) **N.** Basal diameter of penetrating arteries after vehicle injection (x-axis) versus CNO injection (y-axis) showing no significant difference (−7.5±12.6 %, LME p=1 Bonferroni corrected, n=9 mice, 23 vessels). **O.** Population locomotion-triggered averages after vehicle (black) and CNO (red) injection (n=9 mice, 23 arterioles in FL/HL representation). **P.** Basal diameter of penetrating arteries after vehicle injection (x-axis) versus CNO injection (y-axis) showing no significant difference (+7.3±16.4%, LME p=0.81 Bonferroni corrected, n=6 mice, 16 vessels). **Q.** Population locomotion-triggered averages after vehicle (black) and CNO (red) injection (n=6 mice, 16 arterioles in FL/HL representation). Data in **R** and **S** are from penetrating arteries in mice after L-NAME and aCSF infusions. Data in **T** and **U** is from penetrating arteries in mice L-arginine and aCSF infusions. **R.** Basal diameter of penetrating arteries after vehicle infusion (x-axis) versus L-NAME infusion (y-axis). There was no significant change in arterial diameter (−14.8±10.9%, LME p=0.56 Bonferroni corrected, n=6 mice, 14 vessels). **S.** Population locomotion-triggered averages after vehicle (black) and L-NAME (red) infusion (n=6 mice, 14 arterioles in FL/HL representation). **T.** Basal diameter of penetrating arteries after vehicle infusion (x-axis) versus L-arginine infusion (y-axis) showing no significant difference (3.1±16.2%, LME p=0.77 Bonferroni corrected, n=5 mice, 11 vessels). **U.** Population locomotion-triggered averages after vehicle (black) and L-arginine (red) infusion (n=5 mice, 11 arterioles in FL/HL representation). Data in **V** and **W** are from penetrating arteries in mice after CP-99994 and aCSF infusions. Data in **X** and **Y** is from penetrating arteries in mice after Substance P and aCSF infusions. **V.** Basal arteriole diameter of penetrating arteries after vehicle infusion (x-axis) versus CP-99994 infusion (y-axis) showing no significant difference (5.4±23.6%, LME p=1 Bonferroni corrected, n=6 mice, 11 vessels). **W.** Population locomotion-triggered averages after vehicle (black) and CP-99994 (red) infusion (n=6 mice, 11 arterioles in FL/HL representation). **X.** Basal arteriole diameter of penetrating arteries after vehicle infusion (x-axis) versus Substance P infusion (y-axis) showing diameter was significantly increased by substance P (23.3±22.3%, LME p=1.1×10^−2^ Bonferroni corrected, n=6 mice, 13 vessels). **Y.** Population locomotion-triggered averages after vehicle (black) and Substance P (red) infusion (n=6 mice, 13 arterioles in FL/HL representation).

**Supplementary Figure 3.**
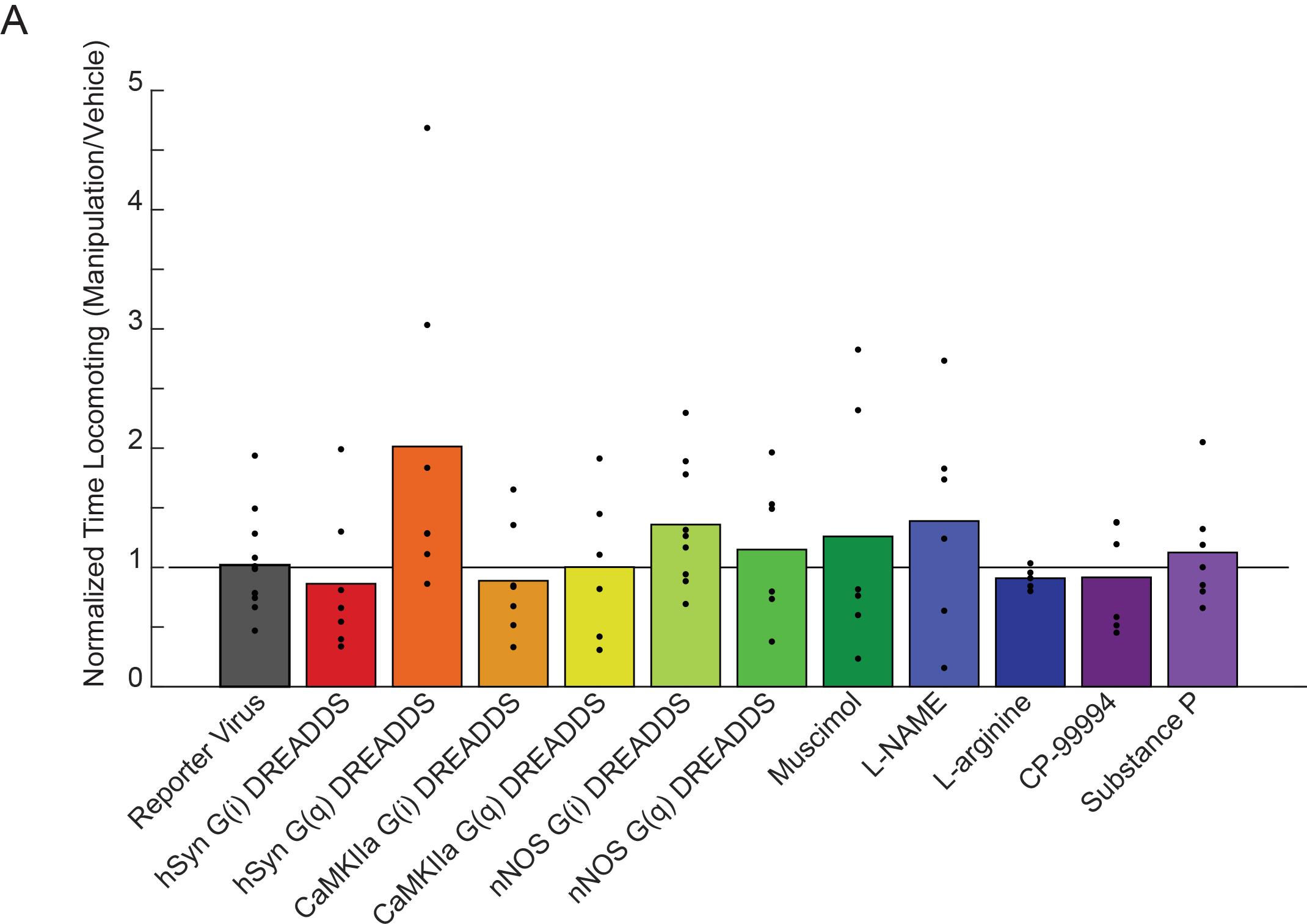
Effects of chemogenetic and pharmacological manipulation on locomotion behavior. Plot showing the amount of time locomoting during 2PLSM imaging for each manipulation normalized by the amount of time locomoting with the vehicle. No manipulation resulted in a significant change in the behavior (LME, all p-values were Bonferroni corrected by the number of different viruses used or by the number of different drugs infused: *Reporter Virus* p=1 n=12 mice; *muscimol* p=1 n=6 mice; *hSyn-G(i) DREADDs* p=1 n=7 mice; *hSyn-G(q) DREADDs* p=0.3 n=7 mice; *CaMKIIa-G(i) DREADDs* p=1 n=7 mice; *CaMKIIa-G(q) DREADDs* p=1 n=6 mice; *nNOS-G(i) DREADDs* p=0.2 n=9 mice; *nNOS-G(q) DREADDs* p=1 n=6 mice; *L-NAME* p=1 n=6 mice; *L-arginine* p=1 n=5 mice; *CP-99994* p=1 n=7 mice; *Substance P* p=1 n=7 mice).

**Supplementary Figure 4.**
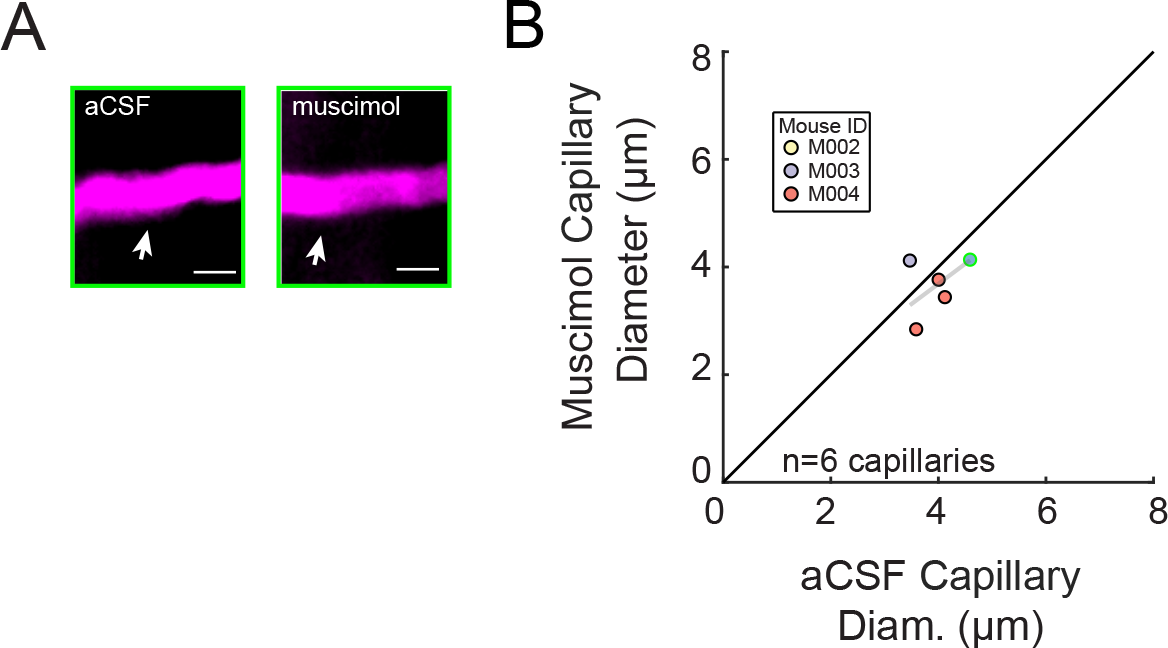
Muscimol infusion does not significantly affect capillary diameter. **A.** Representative images of the same capillary after vehicle infusion (left) and after muscimol infusion (right). There was no significant change seen after muscimol infusion (scale bar 5μm). **B.** Capillary diameter measured after vehicle infusion (x-axis) vs. after muscimol infusion (y-axis) showed no significant changes (0.12±1.1μm LME p=0.66, n=3 mice, 5 capillaries). The point outlined in green is representative image from **A**.

**Supplementary Figure 5.**
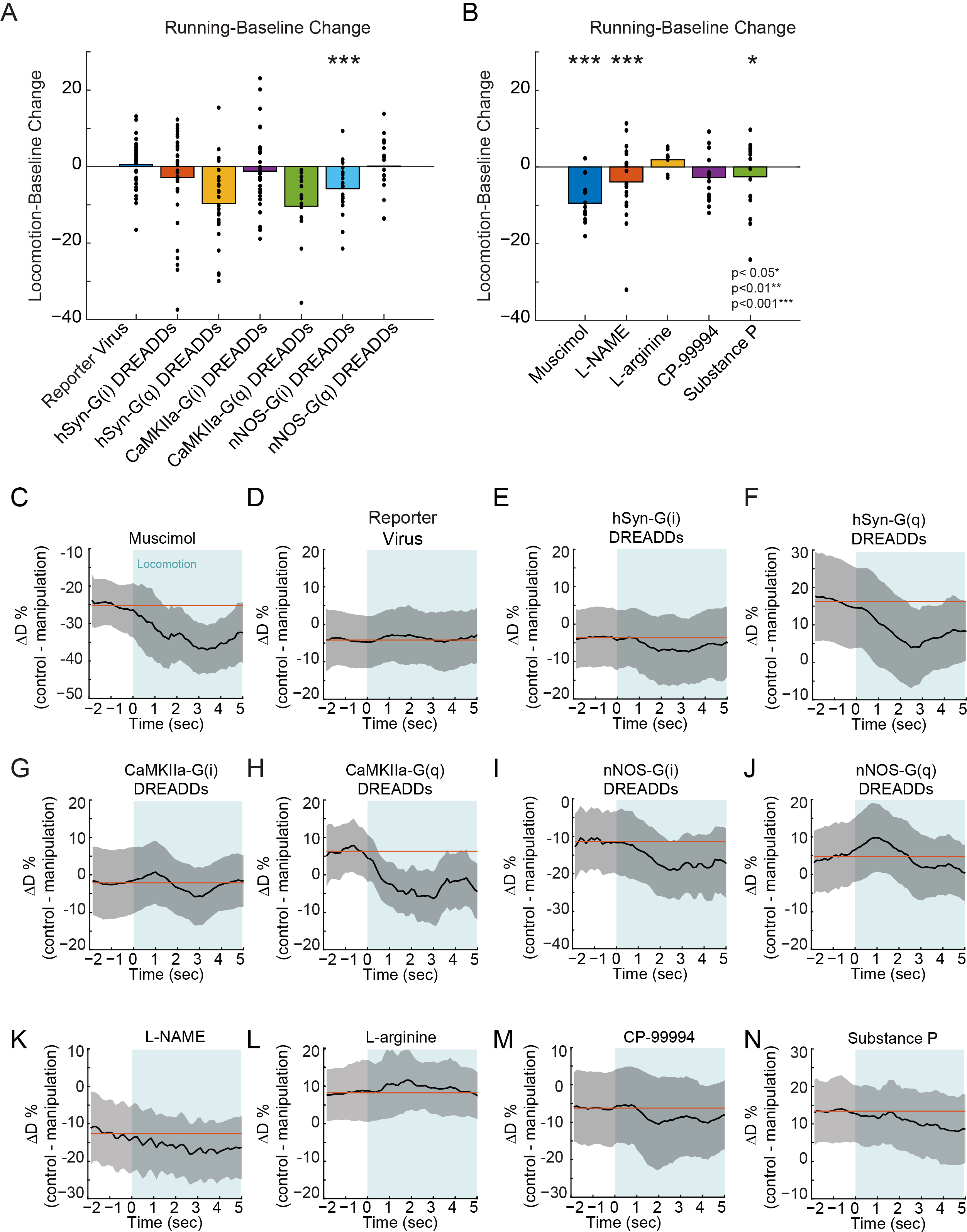
The effects of chemogenetic and pharmacological infusions on evoked arterial diameter changes. The locomotion-baseline changes are calculated from the locomotion fractional difference values (in C-N). The average from −2 to 0 seconds before locomotion onset was subtracted from the average locomotion period, 1 to 4 seconds after locomotion onset. **A.** Locomotion-baseline change for all the virus injected groups. (All p-values, paired t-test, Bonferroni corrected, reporter virus p=1 n=12 mice; hSyn-G(i) DREADDs p=1 n=7 mice; hSyn-G(q) DREADDs p=0.55 n=7 mice; CaMKIIa-G(i) DREADDs p=1 n=7 mice; CaMKIIa-G(q) DREADDs p=1 n=6 mice; nNOS-G(i) DREADDs p=5.2×10^−4^ n=9 mice; nNOS-G(q) DREADDs p=1 n=6 mice) **B.** Locomotion-baseline change for all the infusion groups. (All p-values, paired t-test, Bonferroni corrected, muscimol p=1.6×10^−7^ n=6 mice; L-NAME p=1.2×10^−4^ n=6 mice; L-arginine p=0.49 n=5 mice; CP-99994 p=1 n=7 mice; Substance P p=0.04 n=7 mice) Locomotion-triggered averages (LTA) were calculated by taking any locomotion event >5 seconds in duration with at least 2 seconds of no locomotion before the event. These averages were normalized by the average basal diameter of the same vessel under vehicle control conditions. The locomotion fractional difference was calculated by subtracting the treatment LTA from the vehicle LTA. **C-N** are the individual locomotion fractional diameter differences for each manipulation. The red line shows the mean diameter difference averaged over the two seconds before locomotion onset. A decrease in the values during locomotion relative to the pre-locomotion baseline means that the treatment blocked or (in the case of hSyn G(q) coupled DREADDs) occluded the dilation, which is consistent with the process being involved in evoked dilations

**Supplementary Figure 6.**
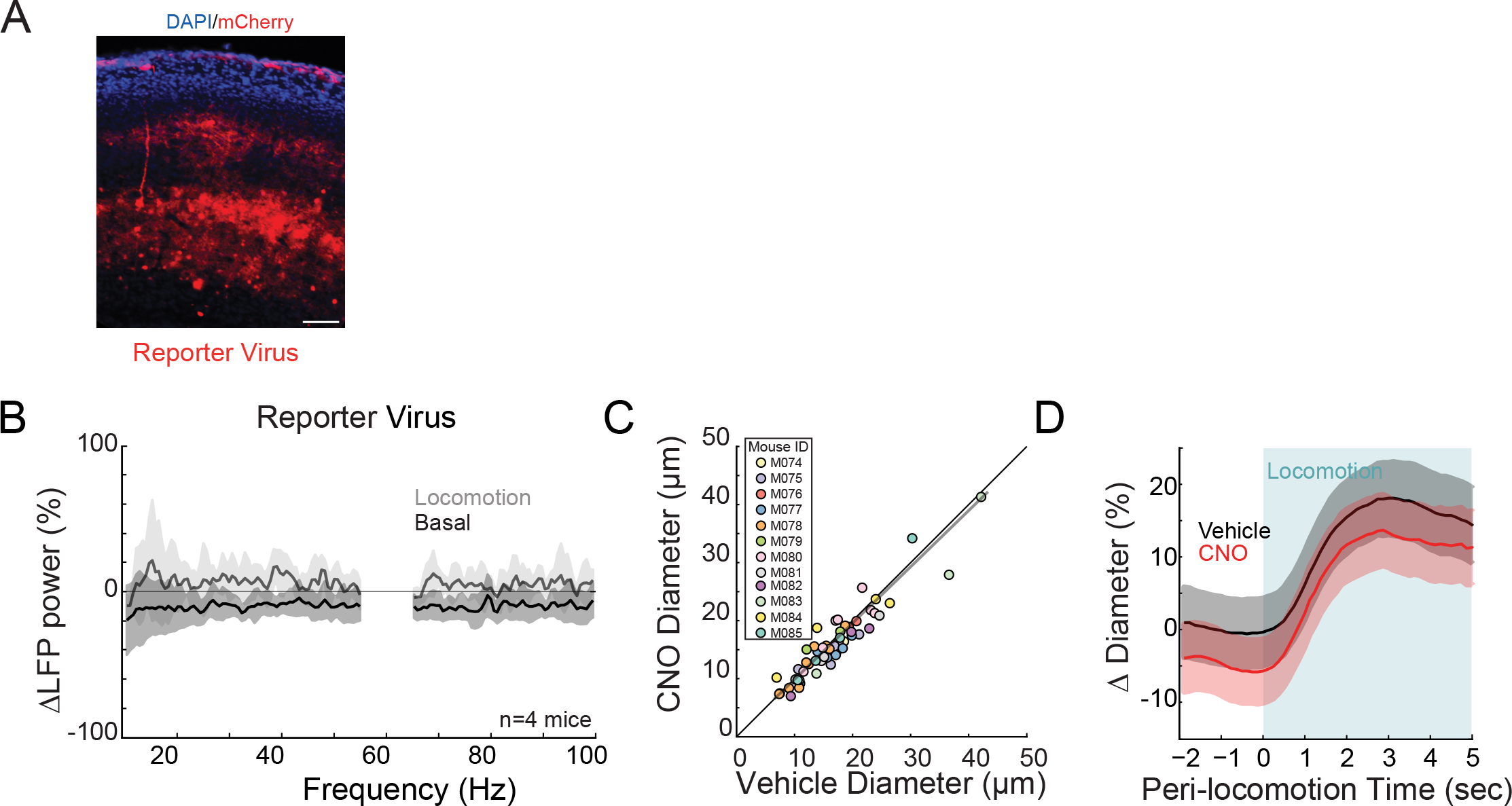
No significant effect of CNO on basal vessel diameter or neural activity. **A.** Representative 90μm thick coronal section of the cortical region infected with the reporter virus showing mCherry expressing neurons (red) and cell nuclei (DAPI, blue). Scale bar is 100μm. Data in **B-D** are from mice injected with AAV-CMV-TurboRFP-WPRE-rBG (fluorescent reporter virus) **B.** LFP power spectra during stationary periods (basal) and locomotion after CNO injection, normalized to vehicle injection in the same mouse. The fluorescent reporter virus does not significantly change neural activity in the gamma band (basal, −9.7±10.6%, paired t-test p=0.17; locomotion, 5.4±8.8 %, paired t-test p=0.29, n=4). Shading denotes mean ± standard deviation. **C.** Plot of basal arteriole diameter after vehicle injection (x-axis) versus CNO injection (y-axis). There were no significant changes in basal diameter (−3.2±14.5%, LME p=0.46 Bonferroni corrected, n= 12 mice, 56 vessels). **D.** Locomotion-triggered averages after vehicle (black) and CNO (red) injections (n= 12 mice, 39 arterioles in FL/HL representation). For both cases, the diameters were normalized by the average basal diameter of the vessel after vehicle injection.

**Supplementary Figure 7.**
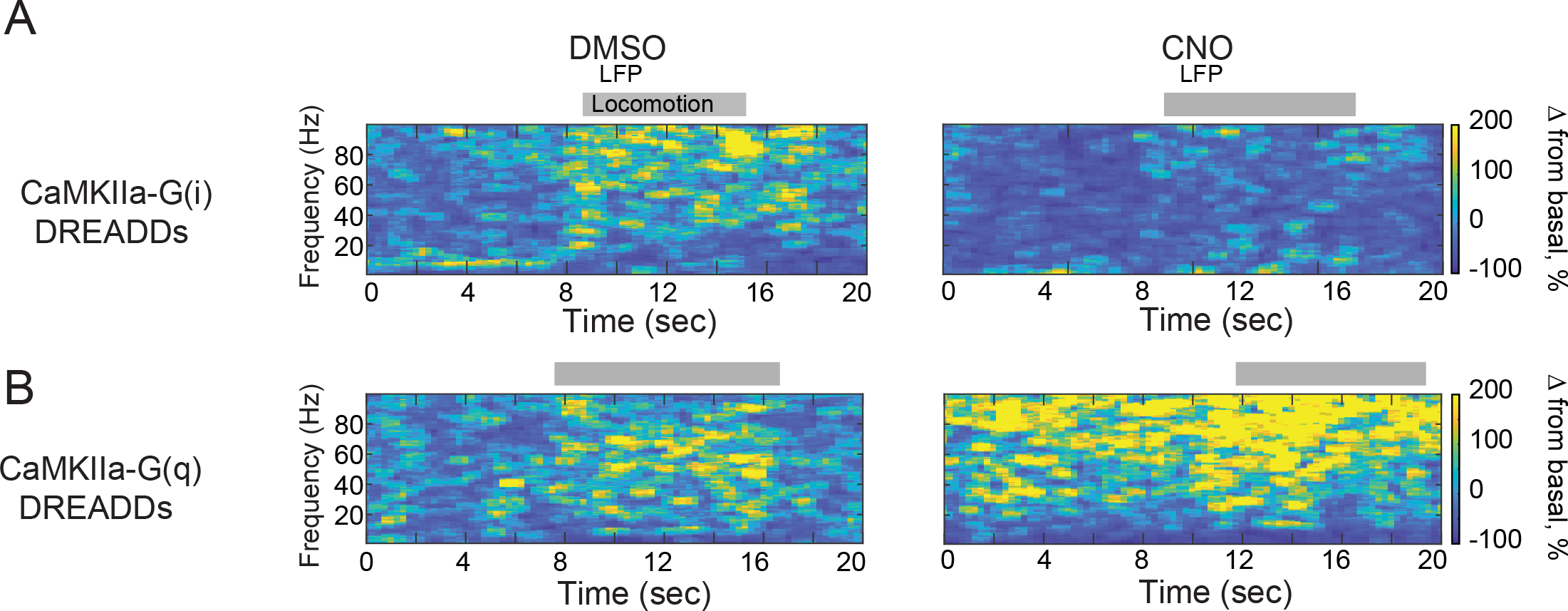
Expressing DREADDs in excitatory neurons produced large changes in overall neural activity. **A.** Representative LFP spectrograms of CaMKIIa-G(i) DREADD infected mouse after vehicle (left) or CNO (right) injection. Locomotion events are denoted by shading. Note the large decrease in neural activity after CNO injection. **B.** Representative LFP spectrograms of CaMKIIa-G(q) DREADD infected mouse (expression in pyramidal neurons) after vihicle (left) or CNO (right) injection. Locomotion events are denoted by shading. Note the large increase in neural activity after CNO injection.

**Supplementary Figure 8.**
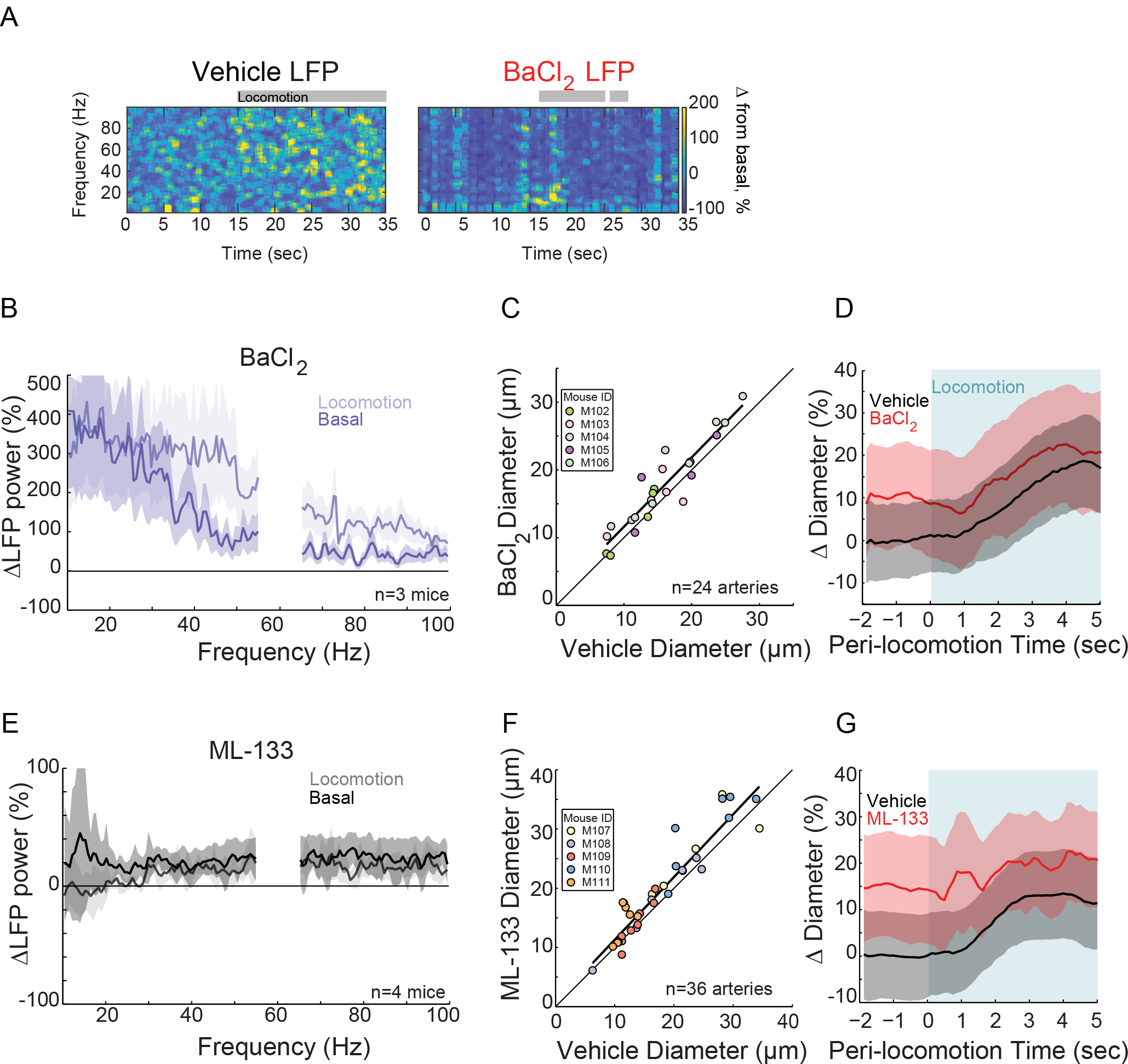
Kir-channel blockers cause large increases in neural activity and basal arterial diameter. **A.** Spectrogram of the LFP in a mouse after infusion with vehicle (left) or BaCl2 (right). The bursts across all frequency bands of the LFP indicate that the BaCl2 infusion induced epileptic-like activity. Locomotion events are denoted with shading. **B.** LFP power spectra during stationary periods (basal) and locomotion after BaCl2 infusion, normalized to vehicle infusion in the same mouse. BaCl2 infusion greatly increases neural activity in the gamma band (basal, +226±188%, paired t-test p=1.1×10^−2^; locomotion, +290±180.1 %, paired t-test p=4.6×10^−2^, n=3). Shading denotes mean ± standard deviation. **C.** Plot of basal arteriole diameter after vehicle infusion (x-axis) versus BaCl2 infusion (y-axis). BaCl2 significantly increased arterial diameter (+12.4±17.2%, LME p=6.9×10^−3^ Bonferroni corrected, n= 5 mice, 24 vessels) **D.** Population locomotion-triggered average after vehicle (black) and BaCl2 (red) infusions (n=5 mice, 16 arterioles in FL/HL representation). For both cases, the diameters were normalized by the average basal diameter of the vessel after vehicle infusion **E.** LFP power spectra during stationary periods (basal) and locomotion after ML-133 infusion, normalized to vehicle infusion in the same mouse (basal, +23.8±3.3%, paired t-test p=0.16; locomotion, +8.7±11.3 %, paired t-test p=0.16, n=4). **F.** Plot of basal arteriole diameter after vehicle infusion (x-axis) versus ML-133 infusion (y-axis). ML-133 significantly increased the basal arterial diameter (+11.8±17.0% LME p=1.3×10^−3^ Bonferroni corrected, n= 5 mice, 36 vessels) **G.** Locomotion-triggered averages after vehicle (black) and ML-133 (red) infusions (n=5 mice, 26 arterioles in FL/HL representation). For both cases, the diameters were normalized by the average basal diameter of the vessel after vehicle infusion

**Supplementary Figure 9.**
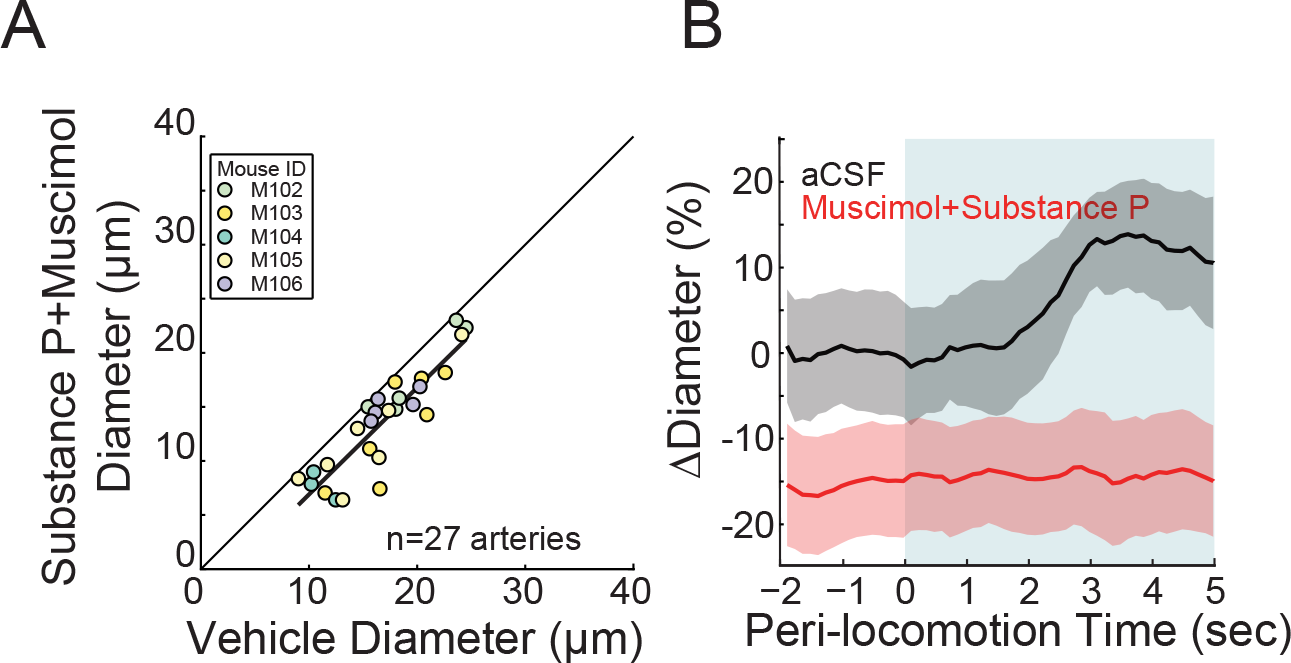
Dilatory effects of Substance P are mediated through local neural excitation, not direct vascular effects. **A.** Plot of basal arteriole diameter after vehicle infusion (x-axis) versus muscimol and substance P infusion (y-axis). The co-infusion of muscimol and Substance P resulted in a significant decrease in basal arterial diameter (−20.0±14.9%, LME p=4.0×10^−7^ Bonferroni corrected, n= 5 mice, 27 vessels) **B.** Locomotion-triggered averages after vehicle (black) and muscimol and substance P (red) infusions (n=5 mice, 16 arterioles in FL/HL representation). For both cases, the diameters were normalized by the average basal diameter of the vessel after vehicle injection. Shading denotes mean ± standard deviation.

**Supplementary Figure 10.**
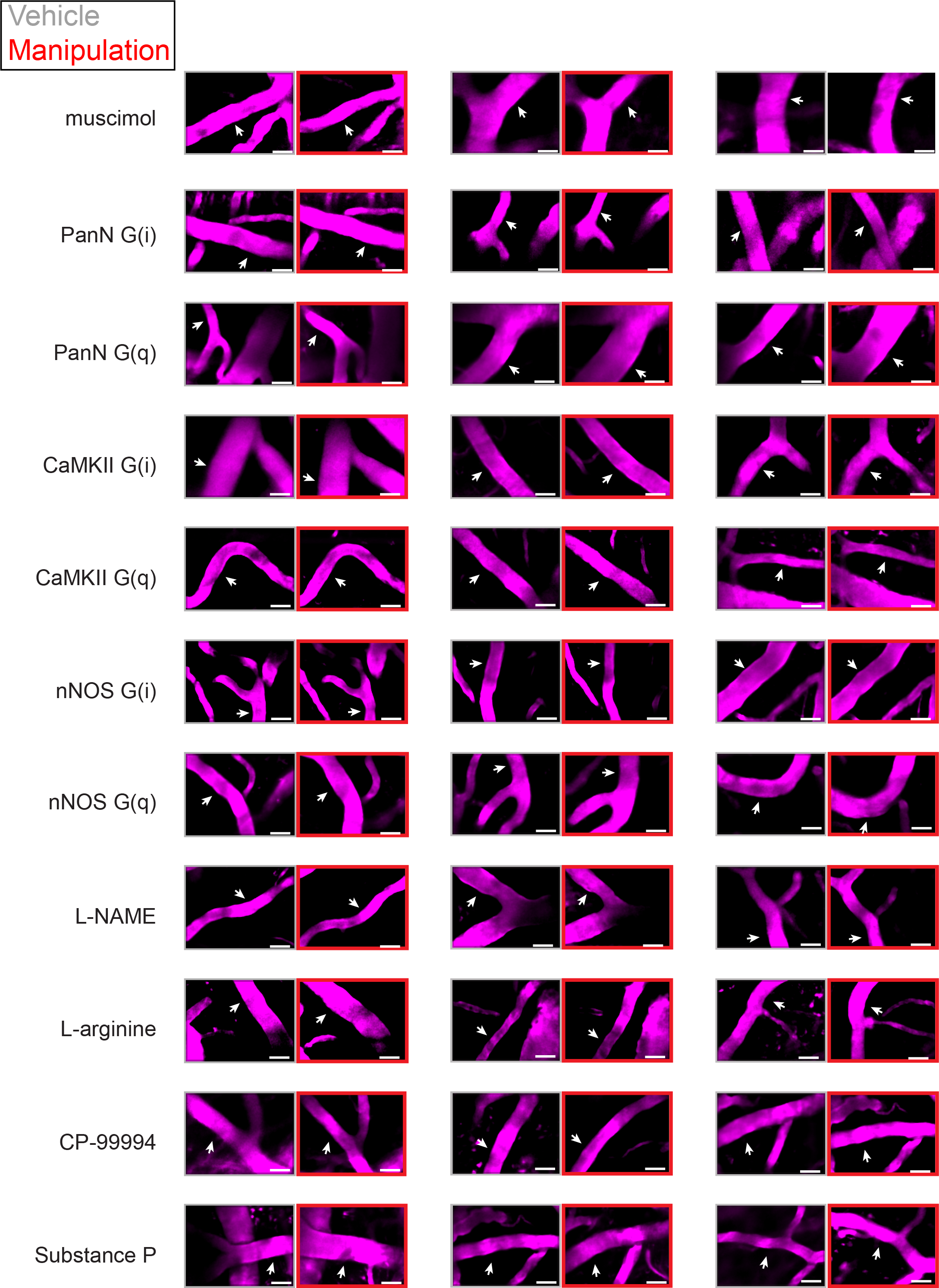
Examples of chemogenetic and pharmacological effects on arteriole diameters. **A.** A maximum projection of two-photon images of arterioles (scale bar 30μm) taken after vehicle treatment (gray box) and chemogenetic/pharmacological effect. The white arrow shows region of diameter measurements

**Supplementary Movie 1**

Locomotion produces a rapid dilation in pial arterioles. This movie shows *in vivo* imaging of a surface arteriole diameter in the somatosensory cortex through the PoRTs window using two-photon microscopy. Left, behavioral camera. Right, FITC-filled arteriole diameter (region marked by white arrow) was plotted versus time in green.

**Supplementary Movie 2**

